# ConvNTC: Convolutional neural tensor completion for predicting the disease-related miRNA pairs and cell-related drug pairs

**DOI:** 10.1101/2024.10.21.619432

**Authors:** Pei Liu, Xiao Liang, Yue Li, Jiawei Luo

## Abstract

Systematic investigation of high-order molecular interactions can deepen our understanding of the mechanisms underlying biological systems. However, effectively capturing both multilinear and nonlinear relationships to accurately identify the complex triplet relationships remains a challenge. In this paper, we present a novel **Conv**olutinal **N**eural **T**ensor **C**ompletion (ConvNTC) to model triplet network connections. ConvNTC consists of a multilinear module and nonlinear module. The former is a tensor decomposition approach that integrates multiple constraints to learn the tensor factor embeddings. The latter contains three components: an embedding generator to produce position-specific index embeddings for each tensor entry in addition to the factor embeddings, a convolutional encoder to perform nonlinear feature mapping while preserving the tensor’s rank-one property, and a Kolmogorov-Arnold Network (KAN) based predictor to effectively capture high-dimensional relationships aligned with the intrinsic structure of real-world data. We demonstrate ConvNTC on two triplet prediction tasks of the disease-related miRNA pairs and cell-related drug pairs, respectively. Comprehensive experiments against ten state-of-the-art methods demonstrate the superiority of ConvNTC in terms of triplet imputation. ConvNTC reveals promising prognostic values of the miRNA-miRNA interactions on breast cancer and detects synergistic drug combinations in cancer cell lines. The source code is available at https://github.com/Liangyushi/ConvNTC.

## Introduction

Investigations of synergistic states between biomolecules aim to reveal the interactions and regulatory mechanisms among various molecules within biological systems, which contribute to gaining a deeper insight into cellular functions, signal transduction and metabolic pathways. Among them, the exploration of synergy between miRNAs related to diseases primarily investigates the role of miRNA pairs in disease progression and development, concentrating on indentifying biomarkers at the gene regulation level (Lai et al., 2019; Xu et al., 2011). In contrast, the discovery of drugs combinations in diverse cellular/disease contexts examines the role of drug pairs in enhancing treatment efficacy, focusing on clinical applications and therapeutic outcomes (Cokol et al., 2011). Translating the identification of disease-related miRNA pairs and disease/cell line-associated drug pairs into the prediction task of miRNA-miRNA-disease and drug-drug-cell/disease triple relationships is an effective strategy.

Consequently, by reformulating the prediction of miRNA-miRNA-disease and drug-drug-cell/disease triple relationships as a third-order tensor completion problem, numerous tensor-based computational models have bee developed. For instance, Chen and Li (2018) presented a tensor completion method, called DrugCom, for capturing disease-related drug combinations using representation learning to integrate multiple additional information of drugs and diseases. Liu et al. (2020) integrated multi-view information on miRNAs and diseases to identify potential miRNA-miRNA-disease associations through a tensor completion framework known as miRCom. Although these methods are effective in identifying cooperative miRNA/drug pairs, they are limited to capturing only the multilinear structure of data, while real-world data often exhibit more complex properties, such as nonlinearity rather than just multilinearity.

Subsequently, deep learning algorithms were introduced to effectively capture nonlinear relationships. For example, Luo et al. (2021) proposed a graph attention mechanism-based neural tensor factorization framework, named GraphTF, to capture disease-related miRNA pairs. Although GraphTF can capture nonlinear factor embeddings, it fails to fully represent the nonlinear relationships within a tensor due to its reliance on the Kronecker product for reconstructing the entire tensor. Sun et al. (2020) proposed a deep tensor factorization model, named DTF, which combines a classical tensor decomposition method CP WOPT with multilayer perceptrons (MLPs) to predict the synergy state of drug pairs. Similarly, Han et al. (2024) proposed a novel constrained tensor factorization (CTF) integrated with a deep neural network, termed CTF-DDI, to identify interactions between drugs across different types. However, both DTF and CTF-DDI tend to overlook the low-rank structure of the tensor and may overfit to sparse training data, as they simply feed the learned low-rank factors from tensor decomposition into a MLP.

Recently, KAN (Liu et al., 2024) integrates Kolmogorov-Arnold mathematical theory with neural network architectures to improve the capacity and interpretability of neural networks. Unlike traditional MLP, KAN replaces linear weight matrices with learnable B-spline functions, aiming to efficiently learn and approximate complex high-dimensional functions by decomposing them into a series of simpler functions. KAN and its various variants have been widely applied across diverse scientific fields (Drokin, 2024; Li et al., 2024). However, to our best knowledge, KAN has not been effectively utilized in tensor decomposition.

In this study, we propose a hybrid deep tensor completion model, termed ConvNTC, which collaboratively captures both multilinear and nonlinear relationships to identify disease-related miRNA pairs and cell line-related drug pairs. Given the intricate and high-order dependencies inherent within a three-order tensor, two separate modules are designed for learning multilinear and nonlinear relationships, respectively. In the multilinear module, we incorporate similarity constraints, Hessian regularization and *L*_2,1_ regularization into a traditional tensor decomposition framework to restrict the factors learning. In contrast, the nonlinear module is designed to capture latent high-order nonlinear interactions and enhance predictive performance, which combines convolutional neural networks (CNNs) with FastKAN (a variants of KAN). This module first employs an embedding generator to obtain initial embeddings for various entities, which are subsequently processed by a convolutional encoder and a FastKAN predictor, yielding predicting scores. Extensive comparative experiments against ten benchmark methods demonstrate that ConvNTC exhibits notable superiority, robustness and scalability. Additionally, case studies further illustrate the effectiveness of ConvNTC in identifying novel disease-related miRNA pairs and cell-related drug combinations.

## Materials and methods

ConvNTC contains three parts: dataset module, multilinear relationship learning module and nonlinear relationship learning module. Taking ‘disease-related miRNA pairs’ as an example, the general outline of ConvNTC is shown in Fig. 1.

**Fig. 1.**
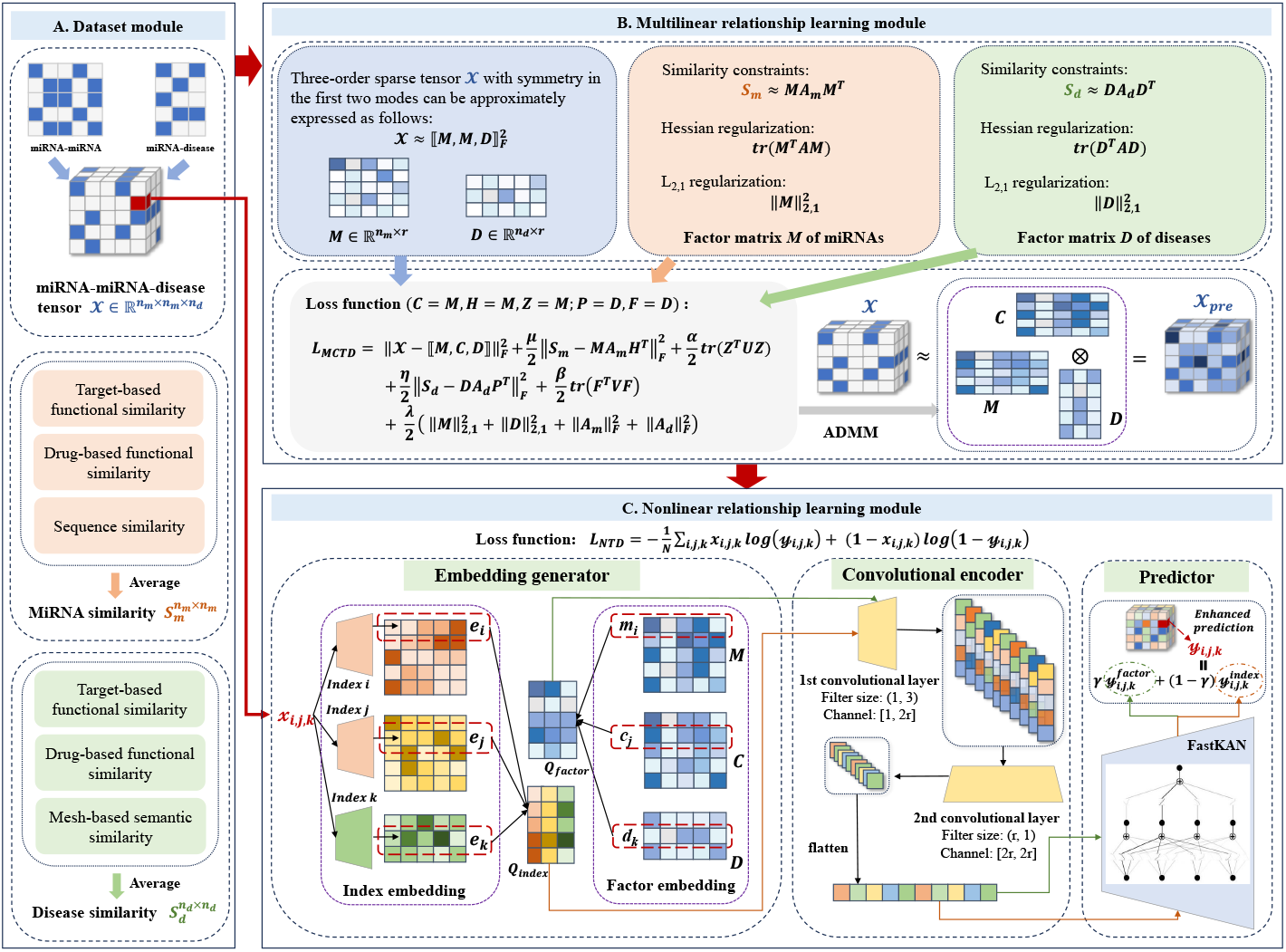
The general outline of ConvNTC by taking the predicting task of disease-related miRNA pairs as an example. It consists of three modules: **A. Dataset module**. We constructed a third-order tensor 𝒳 from miRNA-disease and miRNA-miRNA associations to represent miRNA-miRNA-disease triplets, and obtained miRNA and disease similarity matrices (*S*_*m*_ and *S*_*d*_) using average operation to integrate multi-type similarities. **B. Multilinear relationship learning module**. We learn the multilinear factor embeddings (*M, C* and *D*) with *r* components across three dimensions of the tensor 𝒳 by introducing three constraints into a traditional tensor decomposition. **C. Nonlinear relationship learning module**. An embedding generator first generates the index and factor embeddings of three dimensions for each given entry 𝒳 _i,*j,k*_. Then, a convolutional encoder consisting of two-layer convolutional network is designed to capture the nonlinear features. Finally, a predictor with one-layer FastKAN is adopted to further learn the high-dimensional relationships and obtain the enhanced prediction 𝒴_*i,j,k*_.

### Dataset

**Dataset1** for predicting the disease-related miRNA pairs, was derived from our previous work (Liu et al., 2020), which contains 14,679 miRNA-miRNA-disease triplet relationships among 351 miRNAs and 325 diseases. Additionally, it includes multi-type miRNA and disease similarity matrices. By applying an averaging operation, we obtained the final miRNA and disease similarity matrix. **Dataset2** for predicting cell-related drug pairs, was collected from the previous study (Preuer et al., 2018), which contains 46,124 drug-drug-cell triples with continuous synergy scores among 38 drugs and 39 cell lines. It also contains the preprocessed features of drugs and cell lines. Based on these features, we employed cosine similarity method to obtain the similarity matrices of drugs and cells. More details of these two dataset are provided in Supplementary material 3.

Next, we take the prediction task of disease-related miRNA pairs as an example to describe details of other modules.

### Problem formulation

Given *n*_*m*_ miRNAs, *n*_*d*_ diseases, and their triple associations (*m*_*i*_, *m*_*j*_, *d*_*k*_), a third-order incomplete tensor 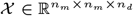 with the first two symmetric modes can be constructed, where *𝒳*_*i,j,k*_ = 1 represents there exists cooperative interaction between miRNA pair (*m*_*i*_, *m*_*j*_) and disease *n*_*d*_, and entry *𝒳*_*i,j,k*_ = 0 represents there is no interaction or missing value, where *i, j* ∈ {1, 2, …, *n*_*m*_}, *k* ∈ {1, 2, …, *n*_*d*_} are the indices of miRNA and disease respectively. Based on this third-order tensor *𝒳* with partially observed interactions between diseases and miRNA pairs, our purpose is to predict whether there exists synergy status of potential miRNA pairs related to a specific disease *d*_*i*_ or not, by filling in the missing values of incomplete third-order tensor *𝒳*.

### Multilinear relationship learning

#### ***M****ulti-****C****onstraint* ***T****ensor* ***D****ecomposition (MCTD)*

Given a third-order miRNA-miRNA-disease incomplete tensor 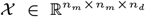 with the first two symmetric modes, and two similarity matrices 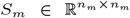 and 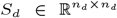 of miRNAs and diseases, we designed a multi-constraint tensor decomposition model, which incorporates similarity constraint, Hessian regularization and *L*_2,1_ regularization into the traditional INDSCAL framework (Carroll and Chang, 1970) to enhance its overall efficacy and provide more effective model constraints. Specifically, similarity constraint can bring in additional biological information on miRNA and disease feature factors. Hessian regularization is more closely aligned with the inherent structure of data compared to conventional Laplace regularization, which can mitigate overfitting after dimensionality reduction. The *L*_2,1_ norm enables to govern the row-wise sparsity, which can prevent overfitting and contribute to a more stable optimization process. The objective function of MCTD is formally defined as follows:

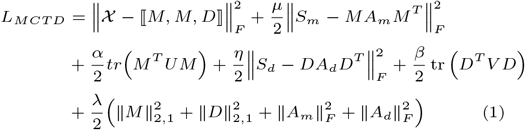

where *r* is the rank of tensor 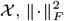 is the Frobenius norm and ⟦· ⟧ is the outer product operation used to reconstruct tensor; 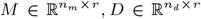 are factor matrices for miRNAs and diseases respectively, while *A*_*m*_, *A*_*d*_ ∈ ℝ^*r×r*^ are their projection matrices; *μ* and *η* are parameters to qualify the contribution of *S*_*m*_ and *S*_*d*_ respectively; 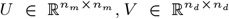 are the Laplace matrices calculated from *S*_*m*_ and *S*_*d*_ respectively, *tr*(·) denotes the Hessian regularization term and *α, β* are parameters to control it; *λ* is a parameter to control *L*_2,1_ regularization.

#### Optimization strategy of MCTD

We employed the ADMM method as optimization strategy of MCTD. To facilitate the solution of *M* and *D*, letting *C* = *M, Z* = *M, H* = *M* and *F* = *D, P* = *D*, the augmented Lagrangian function of MCTD is constructed as follows:

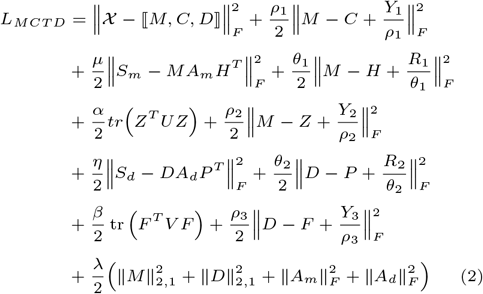

where *Y*_1_, *Y*_2_, 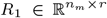, *Y*_3_, 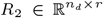 are the Lagrange multipliers; *ρ*_1_, *ρ*_2_, *ρ*_3_, *θ*_1_, *θ*_2_ are the penalty parameters of them. Subsequently, each variable is updated by fixing others until convergence. The updating equations of all variables are provided in Supplementary material 1, and the pseudocode of MCTD is shown in Supplementary algorithm S1. To this end, we can obtain the learned factor matrices *M, C, D* until the above optimization process converges.

### Nonlinear relationship learning

The core of nonlinear relationship learning module is a **N**eural **T**ensor **D**ecomposition (NTD) method, which consists of three components: embedding generator, convolutional encoder and FastKAN predictor. For a third-order incomplete tensor 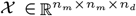 with the first two symmetric modes, the input of NTD is an index vector *t* = [*i, j, k*] and the output is an enhanced prediction score 𝒴_*i,j,k*_.

#### Embedding generator

Given a tensor entry 𝒳_*i,j,k*_ at index (*i, j, k*), the embedding generator takes an index vector *t* = [*i, j, k*] as input, and outputs the corresponding index and factor embeddings *Q*_*index*_, *Q*_*factor*_ of these indices. The index embedding of each indice is produced by a trainable operator *emb*(·) as follows:

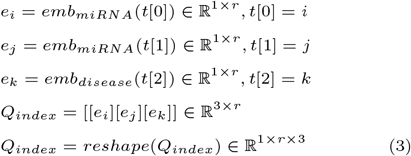

where *emb*(·) is defined as *nn*.*Embedding* in torch, which can map discrete inputs (i.e., miRNA and disease) to a continuous vector. Its factor embedding *Q*_*factor*_ ∈ ℝ^1*×r×*3^ is straightforwardly generated from the learned factors *M, C, D*.

#### Convolutional encoder

The convolutional encoder is composed of two 2D convolution layers: *conv*2*d*_1_(·) with filter size (1, *n*) and *conv*2*d*_2_(·) with a filter size (*r*, 1), where *n* = 3 and *r* are the dimension and rank of association tensor, respectively. Let *n*_*C*_ be the number of channels in the convolutional layers and *δ*_1_, *δ*_2_ are convolution filters of the two convolution layers. It should be emphasized that factor embedding *Q*_*factor*_ shares the convolutional encoder with index embedding *Q*_*index*_. Taking *Q*_*index*_ as an example, its convolutional coding is:

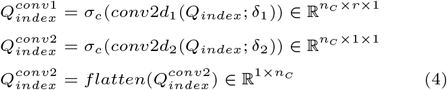

where *σ*_*c*_ = *ReLU* (·) is nonlinear activation function. Similarly, the convolutional coding of *Q*_*factor*_ is 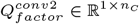.

#### FastKAN predictor

The FastKAN predictor comprises a one-layer of FastKAN (Li, 2024) that utilizes radial basis functions (RBFs) with Gaussian kernels to approximate third-order B-spline bases, thereby significantly accelerating model computations. More explanation are in Supplementary Material 2. Taking index embedding 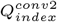 as the input, the output 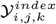 is:

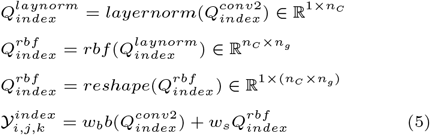

where *rbf* (·) denotes RBFs; *layernorm*(·) is a layer normalization that ensures the input remains within a RBF domain; *b*(·) = *SiLU* (·) is a base activation function similar to residual connection; *w*_*b*_, *w*_*s*_ are trainable factors to better control the overall magnitude of *b*(·) and *rbf* (·) functions. Similarly, the output 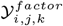 of factor embedding 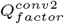 is calculated in the same way, and the final enhanced predicting score is represented as:

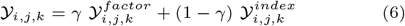

where *γ* is a hyperparameter to measure the weight of index and factor embeddings. To this end, the binary cross-entropy is used to train NTD in a mini-batch way as follows:

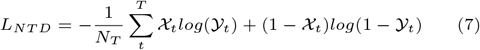

where *N*_*T*_ is the number of training entries in tensor 𝒳, and 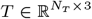 is a training index matrix, and *t* is an index vector of an entry in 𝒳.

## Results

To systematically evaluate the performance of ConvNTC, numerous experiments were conducted for ConvNTC and ten baseline methods (including five linear models, i.e., CANDECOMP/PARAFAC (CP) (Kolda and Bader, 2009), TFAI (Narita et al., 2012), DrugCom (Chen and Li, 2018), TDRC (Huang et al., 2021), CTF (Han et al., 2024), and five nonlinear models, i.e., DeepSynergy (Preuer et al., 2018), Costco (Liu et al., 2019), DTF (Sun et al., 2020), GraphTF (Luo et al., 2021), CTF-DDI (Han et al., 2024)). Details about experimental setup (including dataset division, baseline methods, evaluation metrices and parameter setting) are provided in Supplementary Material 3. Among them, the parameters analysis was conducted to gain the optimal parameter combination of ConvNTC (Details and results are provided in Supplementary Material 4).

### Ablation study

ConvNTC comprises five key components (i.e., MCTD, index embedding, factor embedding, convolutional encoder and FastKAN predictor), affecting on its performance. To assess the impact of these parts, we systematically designed five variants: ConvNTC-nfact, ConvNTC-nind, ConvNTC-nconv, ConvNTC-mlp and ConvNTC-nntd. Summarization is provided in Supplementary Table S3. The results in Supplementary Table S4 depicted that ConvNTC achieved the best performance on both datasets using all five parts. Specifically, we found that the entire nonlinear relationship learning module (i.e., NTD) is able to significantly enhance the performance of ConvNTC. Moreover, convolutional encoder and FastKAN predictor are indispensable parts of NTD, playing a crucial role in capturing the complex nonlinear charateristics. Together, the results clearly demonstrated the advantages of collaborative learning of multilinear and nonlinear modules of ConvNTC. More detailed analysis is in Supplementary Material 5.

### Comparsion with SOTA methods

#### Performance analysis

Table 1 demonstrated that ConvNTC achieved the best results across all metrics on both datasets. Notably, nonlinear methods generally outperformed linear models, highlighting their effectiveness in capturing complex tensor structures. However, most nonlinear approaches struggled to perform consistently well across both datasets, with the exception of DTF. This indicated that incorporating multilinear factor embeddings aided in modeling nonlinear relationships, thereby improving prediction accuracy. ConvNTC’s superior performance over DTF further corroborated the efficacy of its design. The convolutional encoder preserves the tensor’s rank-one structure during nonlinear mapping, and the FastKAN predictor effectively captures complex high-order relationships more closely aligned with tensor’s intrinsic property, resulting in enhanced predictive capability. The p-values of all paired t-test were less than 0.05, further demonstrating that ConvNTC was significantly better than baselines in statistical testing.

**Table 1.**
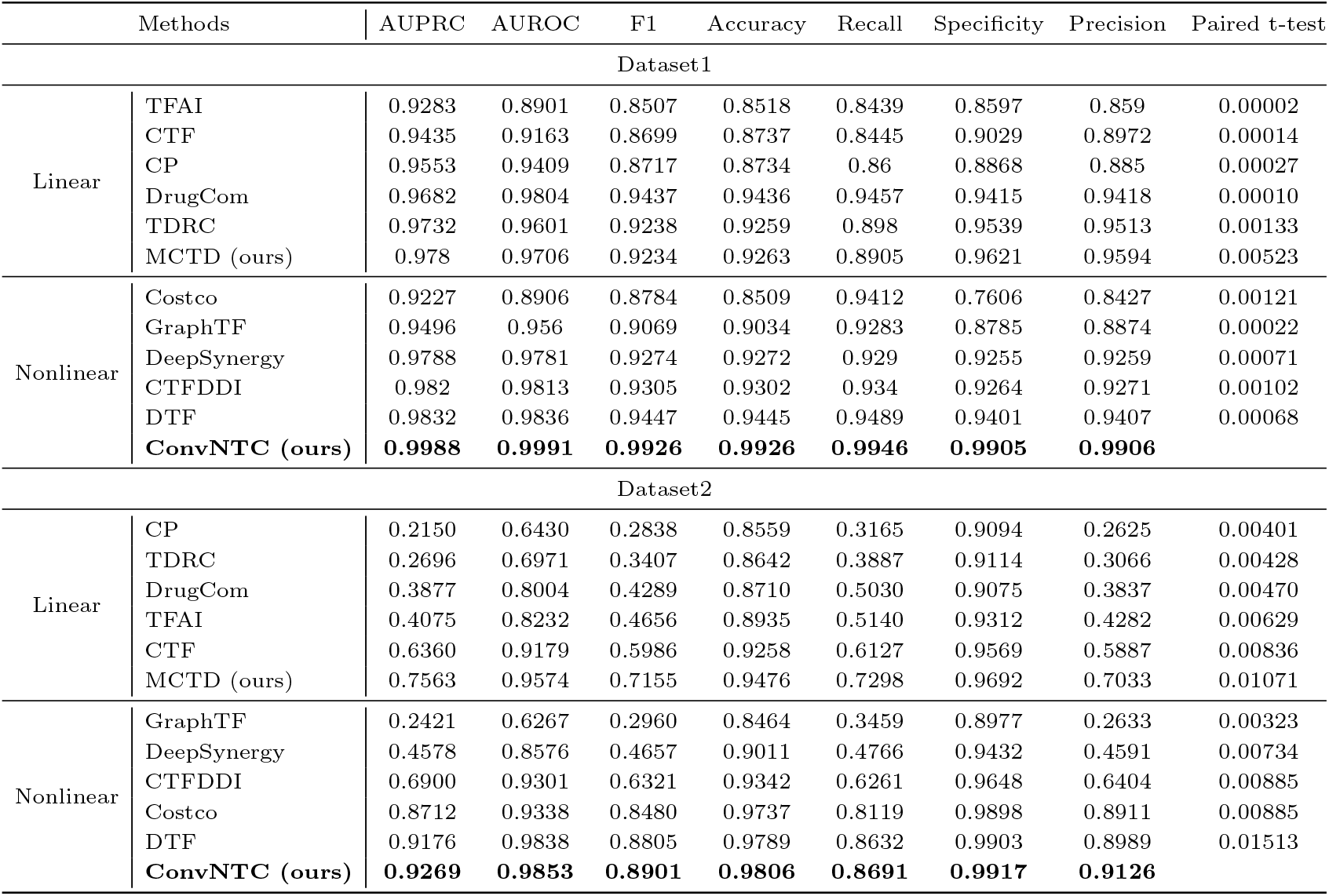
Comparison results of ConvNTC and baselines on Dataset1 and Dataset2. Bolded items indicate the optimal performance.

#### Robustness analysis

To systematically assess the robustness of ConvNTC, we explored the influence of the negative sample size and training ratio on Dataset1. We first randomly sampled the negative samples with a size of [1*n*, 2*n*, 4*n*, 6*n*, 8*n*, 10*n*], where *n* is the number of positive samples. Supplementary Figure S2 depicted the AUPRC values of ConvNTC and baselines, which indicated that all methods decreased as the increasing of negative sample sizes, and ConvNTC performed best. Additionally, the decrease in ConvNTC was significantly lower than others, indicating that ConvNTC exhibited strong robustness against imbalanced data. To assess the model performance under different training ratio, we set the training ratio as 10%, 30%, 50%, 70%, 90% and 100% of the original known data. Supplementary Figure S3 showed that the AUROC values of most models increase with higher training ratios and ConvNTC consistently outperformed all other baselines, which underscored that ConvNTC exhibited resilience to variations in training ratio.

#### Generalizability analysis

To assess the generalizability of ConvNTC, we designed a new experiment to investigate whether ConvNTC can predict synergy status for new unseen data. Specifically, we conducted five-fold cross-validation in five times on the entities (i.e., removing all triplets involving the entity from the training data) rather than triple associations. As show in Supplementary Figure S4 and Supplementary Table S5, ConvNTC performed best on Dataset1, while MCTD performed best on Dataset2, followed by ConvNTC. This indicated that our proposed method can effectively predict miRNA/drug pairs of new diseases/cells to a certain extent. Additionally, We hypothesize that the inferior performance of ConvNTC compared to MCTD in Dataset 2 may stem from the 5-fold cross-validation conducted on entity-level data, resulting in significant data loss and feature noise. More analysis is in Supplementary Material 6. Overall, our method has good generalization in predicting new disease/cell-associated miRNA/drug pairs.

### Case studies

#### Predicting novel miRNA-miRNA-disease associations

To demonstrate the capability of ConvNTC to predict novel miRNA-miRNA-disease associations, we carried out case studies on Breast Neoplasms. Given a triple association (*m*_*i*_, *m*_*j*_, *d*_*k*_), we validated it by verifying its pairwise associations i.e., (*m*_*i*_, *m*_*j*_), (*m*_*i*_, *d*_*k*_), (*j, d*_*k*_). Specifically, we confirmed the miRNA-disease associations by querying the existing public databases, i.e., dbDEMC 2.0 (Yang et al., 2017), miRCancer (Xie et al., 2013) and PhenomiR 2.0 (Ruepp et al., 2010) and HMDD v4.0 (Cui et al., 2024), and verified the miRNA-miRNA pairs using a miRNA set enrichment analysis tool TAM 2.0 (Li et al., 2018). More details on TAM 2.0 are provided in Supplementary Material 7.1.1. Table 2 showed that all predicted top-10 miRNA pairs were confirmed, and their association network in Supplementary Figure S5 indicated that most miRNAs act on diseases in the form of pairs (or modules).

**Table 2.** The detailed evidences of the top-10 predicted miRNA pairs for **Breast neoplasms**. Noting that TF, TS and CS are the abbreviations of Transcription Factor, Tissue Specificity and Cell Specificity respectively. The bolded items indicate the max values.

| MiRNA1 | MiRNA2 | MiRNA1-MiRNA2 Evidences |  |  |  |  |  |  | MiRNA1-Disease Evidences | MiRNA2-Disease Evidences | Triplets Evidences |
| --- | --- | --- | --- | --- | --- | --- | --- | --- | --- | --- | --- |
|  |  | Enrichment Analysis using TAM 2.0 (-log10(p-value)) |  |  |  |  |  |  |  |  |  |
|  |  | Function | Disease | TF | Cluster | Family | TS | CS |  |  |  |
| hsa-mir-125a | hsa-mir-99b | 2.39362 | 3.30437 | 3.57840 | 5.70997 | 4.73993 | 3.15366 | 2.95468 | dbDEMC;miRCancer; PhenomiR;HMDDV4 | dbDEMC;PhenomiR; HMDDV4 | 3.25964 |
| hsa-mir-10a | hsa-mir-99b | 0.00000 | 2.82136 | 3.57840 | 0.00000 | 4.73993 | 0.00000 | 0.00000 | dbDEMC;miRCancer; PhenomiR;HMDDV4 | dbDEMC;PhenomiR; HMDDV4 | 0.52288 |
| hsa-mir-19b-1 | hsa-mir-19b-2 | 2.64839 | 3.49975 | 3.73060 | 0.00000 | 5.70997 | 3.86646 | 3.11691 | dbDEMC;PhenomiR | dbDEMC;PhenomiR | 0.12494 |
| hsa-mir-99a | hsa-mir-99b | 3.82391 | 3.67684 | 0.00000 | 0.00000 | 4.73993 | 0.00000 | 0.00000 | dbDEMC;miRCancer; PhenomiR;HMDDV4 | dbDEMC;PhenomiR; HMDDV4 | 1.50864 |
| hsa-mir-19a | hsa-mir-19b-2 | 2.64839 | 3.17420 | 3.73060 | 0.00000 | 5.70997 | 3.86646 | 3.11691 | dbDEMC;miRCancer; PhenomiR;HMDDV4 | dbDEMC;PhenomiR | 0.43180 |
| hsa-mir-181c | hsa-mir-181d | 3.32808 | 2.93417 | 0.00000 | 5.18709 | 5.01100 | 0.00000 | 3.27327 | dbDEMC;PhenomiR; HMDDV4 | dbDEMC;PhenomiR; HMDDV4 | 1.22915 |
| hsa-mir-154 | hsa-mir-494 | 0.00000 | 2.55261 | 0.00000 | 3.25259 | 3.95468 | 0.00000 | 0.00000 | dbDEMC;miRCancer; PhenomiR;HMDDV4 | dbDEMC;miRCancer; PhenomiR;HMDDV4 | 0.21467 |
| hsa-mir-181a-2 | hsa-mir-181d | 3.45204 | 2.92905 | 0.00000 | 0.00000 | 5.01100 | 0.00000 | 0.00000 | dbDEMC;PhenomiR | dbDEMC;PhenomiR; HMDDV4 | 2.16115 |
| hsa-mir-125b-2 | hsa-mir-99b | 2.39362 | 3.20235 | 0.00000 | 0.00000 | 4.73993 | 0.00000 | 0.00000 | dbDEMC;PhenomiR; HMDDV4 | dbDEMC;PhenomiR; HMDDV4 | 1.23657 |
| hsa-mir-100 | hsa-mir-99b | 3.10876 | 3.89752 | 0.00000 | 0.00000 | 4.73993 | 0.00000 | 0.00000 | dbDEMC;miRCancer; PhenomiR;HMDDV4 | dbDEMC;PhenomiR; HMDDV4 | 0.50864 |

To further evaluate the biological significance of triple predictions, we perform survival analysis of breast cancer from The Cancer Genome Atlas (TCGA) on the top-10 miRNA pairs. More details on the execution of survival analysis are in Supplementary Material 7.1.3. Table 2 showed that three miRNA pairs (hsa-mir-125a, hsa-mir-99b), (hsa-mir-99a, hsa-mir-99b) and (hsa-mir-181a-2, hsa-mir-181d) have a significant impact on the patient’s survival probability. Their survival curves and Pearson correlation of miRNA expression were shown in Supplementary Figure S6. For the pair (hsa-mir-125a, hsa-mir-99b), the combined analysis revealed more statistically significant segregation between high-risk and low-risk groups compared to the individual analyses of hsa-mir-125a and hsa-mir-99b. Similar results were obtained for the other two miRNA pairs. These observations indicated that these miRNA pairs have a synergistic effect on breast cancer and exhibit promising prognostic value. Interestingly, the expressions of pairs (hsa-mir-125a, hsa-mir-99b) and (hsa-mir-181a-2, hsa-mir-181d) were positively correlated, whereas the pair (hsa-mir-99a, hsa-mir-99b) was negatively correlated, implying synergistic and antagonistic role of these miRNA pairs, respectively.

Additionally, we performed the enrichment analysis on the common target genes of these three miRNA pairs. More details on the operation of enrichment analysis are provided in Supplementary Material 7.1.4. Figure 2 depicted the enriched terms and pathways that are known to be implicated in the breast cancer progression. Processes like lysosome and vacuole organization support cancer cell survival, while chromosome segregation and mitotic regulation maintain genomic stability. The ‘Breast cancer pathway’ involves hormonal signaling, cell growth and survival mechanisms specific to breast cancer. Other pathways like PI3K/Akt/mTOR promotes tumor growth, and ATM signaling aids DNA repair. Together, these enriched terms and pathways highlight specific metabolic and signaling processes in breast neoplasm.

**Fig. 2.**
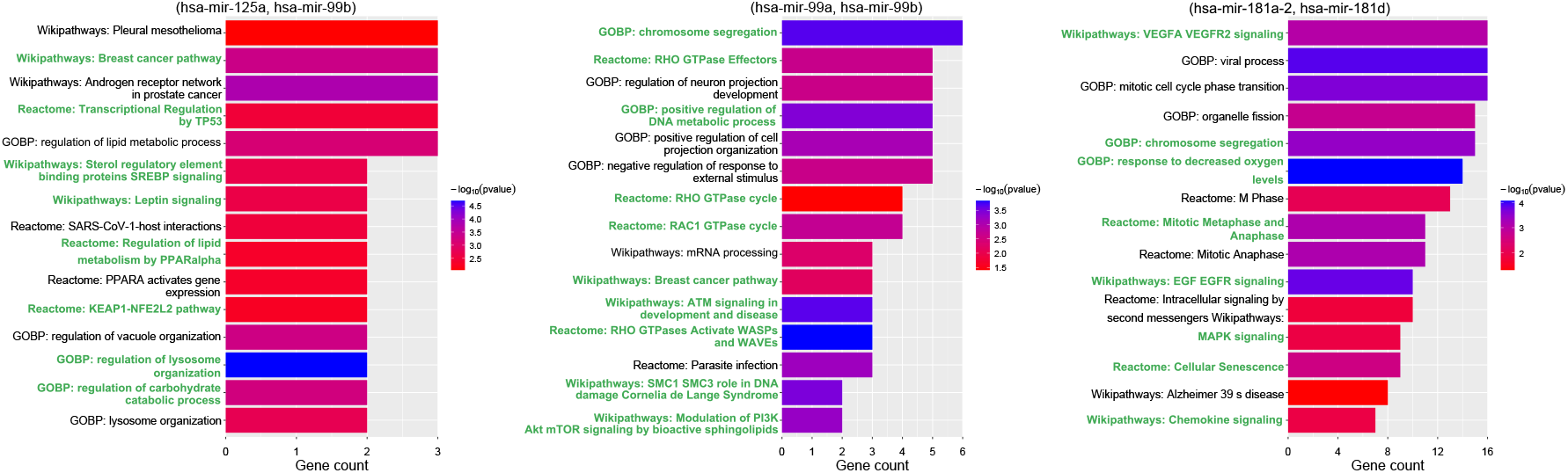
Enrichment analysis of miRNA pairs (hsa-mir-125a, hsa-mir-99b), (hsa-mir-99a, hsa-mir-99b) and (hsa-mir-181a-2, hsa-mir-181d) related to Breast Neoplasms on ‘GO BP’, ‘Wikipathways’ and ‘Reactome’ using R package ‘clusterProfiler’ (Yu et al., 2012) and ‘ReactomePA’ (Yu and He, 2016). Noting that green fonts indicate key processes and pathways related to breast cancer.

#### Predicting novel synergistic drug pairs

We employed ConvNTC to predict novel synergistic drug pairs after training on Dataset2. We showcased the 855 potential drug pairs with 42 drugs for malignant melanoma cancer cell line A375 from paper (Wang et al., 2022). The supports of top-10 predicted drug combinations in Table 3 showed that 9 out of the 10 predicted combinations have been corroborated by prior studies or clinical trials. These combinations focus on inhibiting growth signals, DNA repair mechanisms and cellular metabolism to enhance treatment efficacy for A375 cells. More analysis are provided in Supplementary Material 7.2. Therefore, our drug-drug-cell predictions hold therapeutic potential and warrant further experimental validation.

**Table 3.** The supports of the top-10 predicted novel synergistic combinations on A375 cancer cell line.

| Drug 1 | Drug 2 | Evidences |
| --- | --- | --- |
|  |  | Publications (PMID) |
| Erlotinib | Talazoparib | 36601757; 26124641 |
| Erlotinib | Lenvatinib | 36066408 |
| Erlotinib | Enasidenib | NA |
| Erlotinib | Everolimus | 22378048; 22968184 |
| Erlotinib | Copanlisib | 34734007 |
| Erlotinib | Ruxolitinib | 25065853 |
| Abemaciclib | Erlotinib | 33194700 |
| Erlotinib | Crizotinib | 28961830; 31960422; 31078602; 32540560 |
| Erlotinib | Pazopanib | 29645086; 22688250; 23135778 |
| Erlotinib | Regorafenib | 25907508 |

## Conclusion

By building on the premise of reformulating the task of identifying disease-associated miRNA pairs and cell-related drug pairs as a third-order tensor completion problem, we presented a two-stage hybrid tensor completion framework ConvNTC. Different from prior work, ConvNTC modeled the tensor’s underlying multilinear structure by developing a multi-constraint tensor decomposition method with three constraints. Furthermore, ConvNTC integrated CNN and FastKAN into a unified architecture, which can comprehensively capture the intrinsic high-order complexity of the tensor and the potential nonlinear interactions between objects. As the key contribution, we proposed a convolutional encoder that enabled nonlinear feature mapping while maintaining the tensor’s rank-one property, alongside a FastKAN predictor that effectively captured high-dimensional relationships aligned with the intrinsic structure of the real-world data. This provides a way to enhance model representation capabilities and facilitate feature interaction learning in this domain. Numerous experiments demonstrated the superiority and effectiveness of ConvNTC in predicting triple relationships, and case studies exhibited the significant potential of ConvNTC in advance precision medicine and drug development.

## Acknowledgments

This work is supported by the National Natural Science Foundation of China (#62372165 and #62032007) and the Natural Sciences and Engineering Research Council (NSERC) Discovery Grant (RGPIN-2019-0621).

## Notes

### Competing Interest Statement

The authors have declared no competing interest.

## References

J. D. Carroll and J.-J. Chang. Analysis of individual differences in multidimensional scaling via an n-way generalization of “eckart-young” decomposition. Psychometrika, 35(3):283–319, 1970.

H. Chen and J. Li. Drugcom: Synergistic discovery of drug combinations using tensor decomposition. In 2018 IEEE International Conference on Data Mining (ICDM), pages 899–904. IEEE, 2018.

M. Cokol, H. N. Chua, M. Tasan, B. Mutlu, Z. B. Weinstein, Y. Suzuki, M. E. Nergiz, M. Costanzo, A. Baryshnikova, G. Giaever, et al. Systematic exploration of synergistic drug pairs. Molecular systems biology, 7(1):544, 2011.

C. Cui, B. Zhong, R. Fan, and Q. Cui. Hmdd v4. 0: a database for experimentally supported human microrna-disease associations. Nucleic Acids Research, 52(D1): D1327–D1332, 2024.

I. Drokin. Kolmogorov-arnold convolutions: Design principles and empirical studies. arXiv preprint arXiv:2407.01092, 2024.

G. Han, L. Peng, A. Ding, Y. Zhang, and X. Lin. Ctf-ddi: Constrained tensor factorization for drug–drug interactions prediction. Future Generation Computer Systems, 161:26–34, 2024.

F. Huang, X. Yue, Z. Xiong, Z. Yu, S. Liu, and W. Zhang. Tensor decomposition with relational constraints for predicting multiple types of microrna-disease associations. Briefings in bioinformatics, 22(3):bbaa140, 2021.

T. G. Kolda and B. W. Bader. Tensor decompositions and applications. SIAM review, 51(3):455–500, 2009.

X. Lai, M. Eberhardt, U. Schmitz, and J. Vera. Systems biology-based investigation of cooperating micrornas as monotherapy or adjuvant therapy in cancer. Nucleic acids research, 47(15):7753–7766, 2019.

C. Li, X. Liu, W. Li, C. Wang, H. Liu, and Y. Yuan. U-kan makes strong backbone for medical image segmentation and generation. arXiv preprint arXiv:2406.02918, 2024.

J. Li, X. Han, Y. Wan, S. Zhang, Y. Zhao, R. Fan, Q. Cui, and Y. Zhou. Tam 2.0: tool for microrna set analysis. Nucleic acids research, 46(W1):W180–W185, 2018.

Z. Li. Kolmogorov-arnold networks are radial basis function networks. arXiv preprint arXiv:2405.06721, 2024.

H. Liu, Y. Li, M. Tsang, and Y. Liu. Costco: A neural tensor completion model for sparse tensors. In Proceedings of the 25th ACM SIGKDD International Conference on Knowledge Discovery & Data Mining, pages 324–334, 2019.

P. Liu, J. Luo, and X. Chen. mircom: tensor completion integrating multi-view information to deduce the potential disease-related mirna-mirna pairs. IEEE/ACM Transactions on Computational Biology and Bioinformatics, 19(3):1747–1759, 2020.

Z. Liu, Y. Wang, S. Vaidya, F. Ruehle, J. Halverson, M. Soljačić, T. Y. Hou, and M. Tegmark. Kan: Kolmogorov-arnold networks. arXiv preprint arXiv:2404.19756, 2024.

J. Luo, Z. Lai, C. Shen, P. Liu, and H. Shi. Graph attention mechanism-based deep tensor factorization for predicting disease-associated mirna-mirna pairs. In 2021 IEEE International Conference on Bioinformatics and Biomedicine (BIBM), pages 189–196. IEEE, 2021.

A. Narita, K. Hayashi, R. Tomioka, and H. Kashima. Tensor factorization using auxiliary information. Data Mining and Knowledge Discovery, 25:298–324, 2012.

K. Preuer, R. P. Lewis, S. Hochreiter, A. Bender, K. C. Bulusu, and G. Klambauer. Deepsynergy: predicting anti-cancer drug synergy with deep learning. Bioinformatics, 34(9): 1538–1546, 2018.

Ruepp, A. Kowarsch, D. Schmidl, F. Buggenthin Brauner, I. Dunger, G. Fobo, G. Frishman, C. Montrone, and F. J. Theis. Phenomir: a knowledgebase for microrna expression in diseases and biological processes. Genome biology, 11:1–11, 2010.

Z. Sun, S. Huang, P. Jiang, and P. Hu. Dtf: deep tensor factorization for predicting anticancer drug synergy. Bioinformatics, 36(16):4483–4489, 2020.

J. Wang, X. Liu, S. Shen, L. Deng, and H. Liu. Deepdds: deep graph neural network with attention mechanism to predict synergistic drug combinations. Briefings in Bioinformatics, 23(1):bbab390, 2022.

B. Xie, Q. Ding, H. Han, and D. Wu. mircancer: a microrna– cancer association database constructed by text mining on literature. Bioinformatics, 29(5):638–644, 2013.

J. Xu, C.-X. Li, Y.-S. Li, J.-Y. Lv, Y. Ma, T.-T. Shao, L.-D. Xu, Y.-Y. Wang, L. Du, Y.-P. Zhang, et al. Mirna–mirna synergistic network: construction via co-regulating functional modules and disease mirna topological features. Nucleic acids research, 39(3):825–836, 2011.

Z. Yang, L. Wu, A. Wang, W. Tang, Y. Zhao, H. Zhao, and A. E. Teschendorff. dbdemc 2.0: updated database of differentially expressed mirnas in human cancers. Nucleic acids research, 45(D1):D812–D818, 2017.

G. Yu and Q.-Y. He. Reactomepa: an r/bioconductor package for reactome pathway analysis and visualization. Molecular BioSystems, 12(2):477–479, 2016.

G. Yu, L.-G. Wang, Y. Han, and Q.-Y. He. clusterprofiler: an r package for comparing biological themes among gene clusters. Omics: a journal of integrative biology, 16(5):284–287, 2012.

