## Supplemental Materials for "ConvNTC: Convolutional neural tensor completion for predicting the disease-related miRNA pairs and cell-related drug pairs"

October 22, 2024

### 1 Supplementary material 1: Updating equations and pseudocode of MCTD

The ADMM strategy was adopted to updated each variable of MCTD by fixing others until convergence.

**Updating equations of $M, C, H, Z$ related to miRNAs are:**

$$M = (\mathcal{X}_{(1)}E + \mu S_m H A_m^T + \rho_1 C + \theta_1 H + \rho_2 Z - Y_1 - R_1 - Y_2)$$
$$(E^T E + \mu A_m H^T H A_m^T + \frac{\lambda}{2} W_m I + \rho_1 I + \theta_1 I + \rho_2 I)^{-1} \tag{1}$$
$$C = (\mathcal{X}_{(2)}G + \rho_1 M + Y_1)(G^T G + \rho_1 I)^{-1} \tag{2}$$
$$H = (\mu S_m M A_m + \theta_1 M + R_1)(\mu A_m M^T M A_m + \theta_1 I)^{-1} \tag{3}$$
$$Z = (\rho_2 M + Y_2)(\alpha U + \rho_2 I)^{-1} \tag{4}$$

where $\mathcal{X}_{(1)}$, $\mathcal{X}_{(2)}$ are the mode-1 and mode-2 matricization of tensor $\mathcal{X}$; $E = C \bigotimes D, G = M \bigotimes D$ and $\bigotimes$ is the Khatri–Rao product; $W_m \in \mathbb{R}^{r \times r}$ is diagonal matrix and $W_m(k,k) = \frac{1}{2}\|M(:,k)\|_2$.

**Updating equations of $D, P, F$ related to diseases are:**

$$D = (\mathcal{X}_{(3)}J + \eta S_d P A_d^T + \theta_2 P + \rho_3 F - R_2 - Y_3)$$
$$(J^T J + \eta A_d P^T P A_d^T + \frac{\lambda}{2} W_d I + \theta_2 I + \rho_3 I)^{-1} \tag{5}$$
$$P = (\eta S_d D A_d + \theta_2 D + R_2)(\mu A_d D^T D A_d + \theta_2 I)^{-1} \tag{6}$$
$$F = (\rho_3 D + Y_3)(\beta V + \rho_3 I)^{-1} \tag{7}$$

where $\mathcal{X}_{(3)}$ is the mode-3 matricization, $J = M \bigotimes C$, $W_d \in \mathbb{R}^{r \times r}$ is diagonal matrix and $W_d(k,k) = \frac{1}{2}\|D(:,k)\|_2$.

**Updating equations of Lagrange multipliers $Y_1, Y_2, Y_3, R_1$ and $R_2$ are:**

$$Y_1 = Y_1 + \rho_1(M - C), \rho_1 = \epsilon \rho_1 \tag{8}$$
$$Y_2 = Y_2 + \rho_2(M - Z), \rho_2 = \epsilon \rho_2 \tag{9}$$
$$Y_3 = Y_3 + \rho_3(D - F), \rho_3 = \epsilon \rho_3 \tag{10}$$
$$R_1 = R_1 + \theta_1(M - H), \theta_1 = \epsilon \theta_1 \tag{11}$$
$$R_2 = R_2 + \theta_2(D - P), \theta_2 = \epsilon \theta_2 \tag{12}$$

where $\epsilon = 1.15$ is the updating scale of penalty factor.

**Updating rules of projection matrices $A_m$ and $A_d$.** Taking $A_m$ as an example, the optimization problem of $A_m$ is:

$$L_{A_m} = \frac{\mu}{2}\left\|S_m - MA_mM^T\right\|_F^2 + \frac{\lambda}{2}\|A_m\|_F^2 \tag{13}$$

To effectively update $A_m$, we adopted the conjugate gradient method (CG) and its updating rules $CG(S_m, M, M, \mu, \lambda)$ of k-th iteration is:

$$\begin{aligned} a^{(k)} &= \frac{\|R^{(k)}\|_F^2}{\mu\|MB^{(K)}M^T\|_F^2 + \lambda\|B^{(k)}\|_F^2} \\ R^{(k+1)} &= R^{(k)} - a^{(k)}\left(\mu M^T MB^{(K)}M^T M + \lambda B^{(K)}\right) \\ b^{(k)} &= \frac{\|R^{(k+1)}\|_F^2}{\|R^{(k)}\|_F^2}, B^{(k+1)} = R^{(k+1)} + b^{(k)}B^{(k+)} \end{aligned} \tag{14}$$

where $A_m^{(0)} = 0, R^{(0)} = \mu M^T S_m M - \mu M^T M A_m^{(0)} M^T M - \lambda A_m^{(0)}, B^{(0)} = R^{(0)}$.

**Updating tensor $\mathcal{X}$.** Assuming that $O$ is an indicator tensor with the value of 1 and 0 to denote the corresponding position of known and unknown entries in $\mathcal{X}$, the updating formula is:

$$\begin{aligned} \mathcal{X}_{pre} &= [\![M, C, D]\!] \\ \hat{\mathcal{X}} &= \mathcal{X} + (1 - O)\mathcal{X}_{pre} \end{aligned} \tag{15}$$

**The pseudocode of MCTD as follows:**

**Algorithm S1** The pseudocode of MCTD

**Input:** Triple relationship tensor $\mathcal{X}$, similarity matrices $S_m, S_d$, laplace matrices $U, V$ calculated from $S_m, S_d$, and parameters $\mu, \eta, \alpha, \beta, \lambda$

**Output:** Reconstructed tensor $\mathcal{X}_{pre}$, learned factor matrices $M, C, D$

```
Initializing tensor X^0 = X, factor matrices M^0, C^0, D^0 by uniform distribution, other matrices and lagrange multipliers H^0, Z^0, P^0, F^0, Y_1^0, Y_2^0, Y_3^0, R_1^0, R_2^0 = 0, projection matrices A_m^0 = 0, A_d^0 = 0, and parameters ρ_1^0, ρ_2^0, ρ_3^0, θ_1^0, θ_2^0 = 1e − 6, t = 0, t_max = 100, tol = 1e − 4,
while t < t_max do
    Updating A_m^{t+1}, A_d^{t+1} using CG(S_m, M^t, M^t, μ, λ) and CG(S_d, D^t, D^t, η, λ) in Eq.(14)
    Updating M^{t+1}, C^{t+1}, H^{t+1}, Z^{t+1} by Eq.(1-4)
    Updating D^{t+1}, P^{t+1}, F^{t+1} by Eq.(5-7)
    Updating Y_1^{t+1}, Y_2^{t+1}, Y_3^{t+1}, R_1^{t+1}, R_2^{t+1} and ρ_1^{t+1}, ρ_2^{t+1}, ρ_3^{t+1}, θ_1^{t+1}, θ_2^{t+1} by Eq.(8-12)
    Updating X^{t+1} by Eq.(15)
    Calculating err = ||X^{t+1} − X^t||_F^2 / X^t
    if err < tol then
        break
    end if
end while
return X_pre = [[M^{t+1}, C^{t+1}, D^{t+1}]], M^{t+1}, C^{t+1}, D^{t+1}
```

#### 2 Supplementary material 2: Structure of FastKAN

##### 2.1 Kolmogorov-Arnold Network (KAN)

The key idea behind Kolmogorov-Arnold Network (KAN) proposed by Liu et al. (2024), is based on the Kolmogorov-Arnold representation theorem. This theorem allows a multivariate function $f(x)$ to be expressed as a superposition of simpler functions, which can be more easily learned by a neural network. Mathematically, for a smooth $f(x) : [0, 1]^n \to \mathbb{R}$, it can be expressed as:

$$f(x) = \sum_{q}^{2n+1} \Phi_q \Big( \sum_{p}^{n} \phi_{q,p}(x[p]) \Big) \tag{16}$$

where $\Phi, \phi$ are univariate non-linear functions with their own trainable parameters.

For a supervised learning task consisting of input-output pairs $(x[i], y[i])$, KAN aims to find $f$ such that $y[i] \approx f(x[i])$ for all data points. In the original paper, KAN parameterizes each 1D function as a B-spline curve, with learnable coefficients of local B-spline basis functions. A general KAN network is a composition of $L$ layers with shape $[n_0, n_1, ..., n_L]$: given an input vector $\boldsymbol{x_0} = [x[0], x[1], ..., x[n_0]] \in \mathbb{R}^{n_0}$ and the dimension of output $n_L = 1$, the output is:

$$f(x) = \sum_{l_{L-1}=1}^{n_{L-1}} \phi_{L-1,l_L,l_{L-1}} \Big( \sum_{l_{L-2}=1}^{n_{L-2}} ... \Big( \sum_{l_2=1}^{n_2} \phi_{2,l_3,l_2} \Big( \sum_{l_1=1}^{n_1} \phi_{1,l_2,l_1} \Big( \sum_{l_0=1}^{n_0} \phi_{0,l_1,l_0} \Big( x_0[l_0] \Big) \Big) \Big) \Big) ... \Big) \tag{17}$$

where $n_i$ is the number of nodes in the $i$-th layer, $x_0[l_0]$ is the $l_0$-th data point of $\boldsymbol{x_0}$ in the 0-th layer; $\phi_{l,i,j}$ is the activation function connecting the $i$-th neuron in the $(l + 1)$-th layer and the $j$-th neuron in the $l$-th layer. More specifically, for each data point $x$ in vector $\boldsymbol{x}$, the output of activation function $\phi(\cdot)$ in a single KAN layer is:

$$\begin{aligned} \phi(x) &= w_b b(x) + w_s spline(x) \\ b(x) &= SiLU(x) = \frac{x}{1 + e^{(-x)}} \\ spline(x) &= \sum_i c_i B_i(x, n_g, k) \end{aligned} \tag{18}$$

where $b(\cdot)$ is the basis function similar to residual connections, $spline(\cdot)$ is defined as a linear combination of B-splines basis, and $w_b, w_s$ are trainable factors to better control the overall magnitude of $b(\cdot)$, $spline(\cdot)$ functions; $c_i$ is the learnable coefficient, $B_i(\cdot)$ is B-spline basis function, $n_g$ is the number of spline grids, $k$ is the order of B-spline basis. Original paper sets $n_g = 5, k = 3$, so the number of B-splines basis is $n_g + k = 8$

for each data point. The output of $\phi(\cdot)$ is the sum of the basis function $b(\cdot)$ and the spline function $spline(\cdot)$.

This hierarchical structure enables KANs to model complex functions by breaking them down into simpler components, which can be learned more efficiently by the neural network. However, it may still encounter efficiency bottlenecks due to B-spline functions.

#### 2.2 KAN with Radial Basis Function Networks

FastKAN proposed by Li (2024), a new implementation of KANs that significantly accelerates the model calculation. Specifically, FastKAN employs Gaussian kernel-based radial basis functions (RBFs) to approximate 3rd-order B-spline basis. Furthermore, layer normalization is utilized to ensure the input remains within the RBF domain. These modifications result in a more streamlined implementation of FastKAN, while maintaining accuracy.

The fundamental concept behind radial basis functions (RBFs) is to approximate a target function by combining multiple radially symmetric functions, each centered at distinct points within the input space. The output of an RBF network is expressed as a linear combination of these radial basis functions, weighted by adjustable coefficients. Mathematically, for each data point $x$ in vector $\boldsymbol{x}$, the Gaussian kernel-based RBF network with $n_g$ centers (i.e. each data point has $n_g$ Gaussian kernels) can be represented as:

$$rbf(x) = \sum_{i=1}^{n_g} w_i \varphi\Big(\|x - c_i\|\Big)$$
$$\varphi(r) = exp\Big(-\frac{r^2}{2h^2}\Big) \tag{19}$$

where $w_i$ are the adjustable weights or coefficients of the $i$-th center; $\varphi(\cdot)$ is the radial basis function with Gaussian kernel, which depends on the distance between the input $x$ and a center $c_i$; $r$ is the radial distance, and $h$ is a parameter that controls the width or spread of the function. Therefore, for each data point $x$ in vector $\boldsymbol{x}$, the output of the activation function $\phi(\cdot)$ in a FastKAN layer is:

$$\phi(x) = w_b b(x) + w_s \ rbf\Big(layernorm(x)\Big) \tag{20}$$

where $layernorm(\cdot)$ denotes layer normalization. In this paper, we set $n_g = 8$ as in the original paper.

### 3 Supplementary material 3: Experimental setup

To systematically assess the performance of ConvNTC, we conducted five iterations of five-fold cross-validation experiments on two datasets and compared the results with several state-of-the-art methods.

#### 3.1 Dataset

##### 3.1.1 Summarization

Our approach is primarily applied in two downstream predicting tasks i.e., disease-related miRNA pairs and cell-related drug pairs predictions. The details of data are as follows:

- **Dataset1:** Disease-related miRNA pairs were derived from our previous work (Liu et al., 2020), which contains 14,679 miRNA-miRNA-disease triplet relationships among 351 miRNAs and 325 diseases. Specifically, these triplets were constructed from the experimentally verified human miRNA-disease associations from HMDD v3.2 (Huang et al., 2019) and the miRNA-miRNA interactions based on the miRNA family information from miRbase v22 (Kozomara et al., 2019). The specific formulation equation is as follows:

$$X(m_i, m_j, d_k) = \begin{cases} 1, (m_i, m_j) = 1, (m_i, d_k) = 1 \text{ and } (m_j, d_k) = 1 \\ 0, \text{ otherwise.} \end{cases} \tag{21}$$

  Additionally, this dataset includes three miRNA similarities (i.e., target-based functional similarity, drug-based functional similarity and sequence similarity) and three disease similarities (i.e., target-based functional similarity, drug-based functional similarity and Mesh-based similarity). By applying an averaging operation, we obtained the final miRNA similarity matrix and disease similarity matrix.

- **Dataset2:** Cell-related drug pairs were collected from the previous study (Preuer et al., 2018), which contains 46,124 drug-drug-cell triples with continuous synergy scores among 38 drugs and 39 cell lines. Since we only explored the synergy interaction, not synergy effect, we generated binary labels to represent the synergy status by setting the threshold as 30. If the synergy score of a given drug pair is greater than 30, the synergy status is 1, otherwise it is 0. We treated the entries with synergy status 1 as positive samples, and those with synergy status 0 as negative samples. It also contains the preprocessed features of drugs and cell lines (i.e., 4387 chemical features of drugs and 3984 genomic features of cell lines). Based on these features, we employed the cosine similarity method to obtain the similarity matrices of drugs and cells.

##### 3.1.2 Division

- For Dataset 1, we assumed that all missing entries are negative samples, resulted in the number of negative samples being significantly larger than that of positive samples. To address the imbalance between positive and negative samples, we first randomly selected a quantity of negative samples equal to that of the positive samples from the unobserved entries (except for experiments of negative sample analysis). Subsequently, both positive and negative samples are partitioned into five subsets. For each subset, four parts of the positive and negative samples serve as the training set, while the remaining part is designated as the testing set. This procedure is repeated five times and finally, the average performance is reported. It should be noted that for nonlinear models, we also allocated 10% of the training set as a validation set.

- For Dataset 2, following the binarization of labels, there are 4,162 positive samples and 41,942 negative samples, resulting in a positive-to-negative sample ratio of approximately 1:10. However, we do not conducted random sampling of negative samples; instead, we directly implemented 5-fold cross-validation. This approach enables a more intuitive assessment of the model's performance in the context of imbalanced positive and negative samples. The principles governing the division of the Dataset2 remain consistent with those applied to Dataset1.

#### 3.2 Evaluation metrics

We used eight standard supervised learning metrics to evaluate the performance of our proposed method. These include the Area Under Receiver Operating Characteristic curve (AUROC) and the Area Under Precision-Recall curve (AUPRC), which can quantify the predicting accuracy. In addition, we also utilize the following evaluation metrics to obtain a more comprehensive assessment:

$$\begin{aligned} &Accuracy = \frac{TP + TN}{TP + TN + FP + FN}, \\ &Recall = \frac{TP}{TP + FN}, \quad Precision = \frac{TP}{TP + FP} \\ &F1 = \frac{2 \times Precision \times Recall}{Precision + Recall}, \quad Specificity = \frac{TN}{TN + FP} \end{aligned} \tag{22}$$

where $TP$ and $TN$ represent the number of true positives and true negatives respectively, while $FP$ and $FN$ are false positives and false negatives, respectively. To statistically measure the significant improvement of performance between ConvNTC and baselines, we employed the paired t-test on these metrics at a significance level of 0.05.

#### 3.3 Baseline methods

We conducted a comparative analysis of ConvNTC against the following state-of-the-art models for predicting triple associations in bioinformatics. The baselines can be divided into two categories, i.e., linear models and nonlinear models.

##### 3.3.1 Linear models

- CANDECOMP/PARAFAC (CP) (Kolda and Bader, 2009): A classical tensor factorization model without any auxiliary information which decomposes a tensor as a sum of rank-one tensors via alternating least squares (ALS) rules. The input structure of tensor is $O_1 \times O_2 \times O_3$, where $O_1, O_2, O_3$ are three different objects.

- TFAI (Narita et al., 2012): A variant incorporates auxiliary information into the CP model by introducing graph Laplacian regularizations. The input structure of tensor is $O_1 \times O_2 \times O_3$.

- DrugCom (Chen and Li, 2018): A tensor completion method for capturing disease-related drug combinations using representation learning to integrate multiple sources of additional information about drugs and diseases. The input structure of tensor is $O_1 \times O_1 \times O_3$.

- TDRC (Huang et al., 2021): A tensor decomposition method to predict multi-type miRNA-disease associations by constraining the factor matrices with auxiliary information of miRNAs and diseases. The input structure of tensor is $O_1 \times O_2 \times O_3$.

- CTF (Han et al., 2024): A constrained tensor decomposition model for predicting drug–drug–type triple relationship, incorporating drug similarity and other constraints into the CP model. The input structure of tensor is $O_1 \times O_1 \times O_3$.

The parameters except for $r$, of all baselines are set to the same values as those in original papers. Since the rank $r$ is an intrinsic property of the tensor, we uniformly set $r = 57$ for Dataset1 and $r = 123$ for Dataset2 for all models that require rank $r$.

###### 3.3.2 Nonlinear models

- DeepSynergy (Preuer et al., 2018): A deep learning approach for predicting the synergistic scores of drug combinations related to cell lines, consists of a normalization strategy to account for input data heterogeneity and a Multilayer Perceptron (MLP) with shape $[8182, 4096, 1]$ to model drug synergies. The input is the normalized feature of each known entry in a tensor with shape $O_1 \times O_1 \times O_3$.

- Costco (Liu et al., 2019): A versatile neural tensor completion model for sparse tensors, utilizing the expressive capabilities of convolutional neural networks (CNNs) to capture intricate nonlinear interactions within tensors. Its parameter-sharing strategy effectively maintains the desired low-rank structure. Furthermore, CoSTCo is scalable, as it avoids computationally or memory-intensive operations, such as the Kronecker product. The input is the index of each entry in tensor with with a shape of $O_1 \times O_2 \times O_3$, and the dimension of input features is .

- DTF (Sun et al., 2020): A deep tensor factorization model for predicting the synergy status of drug pairs related to cells, integrates a tensor factorization method CP_WOPT with a deep neural network (DNN). DTF mainly adopts CP_WOPT to generate the factors of each mode in tensor, and concentrates them to form the feature of each entry in tensor with shape $O_1 \times O_1 \times O_3$. Then, it normalizes the entry feature in a same way proposed in DeepSynergy, and feeds the normalized entry feature into MLP with shape $[2048, 1024, 512, 1]$ to calculate the predicted probabilities.

- GraphTF (Luo et al., 2021): A graph attention mechanism-based tensor decomposition method for predicting disease-associated miRNA-miRNA pairs. It adopts graph attention network to capture node features over multi-source biological network, and then uses the learned miRNA and disease features to reconstruct the association tensor via the Kronecker product. The inputs are two miRNA similarity matrices, one disease similarity matrix, and a tensor with a shape of $O_1 \times O_3 \times O_1$.

- CTF-DDI (Han et al., 2024): A novel methods for potential drug–drug interactions prediction combines CTF methods with a MLP layer ($[256, 256, 128, 1]$) to extract nonlinear features. It takes the learned tensor factors from CTF as the inputs, and outputs the enhanced predicted scores.

The parameters of all the above nonlinear methods are set to be consistent with the original paper.

To this end, we utilized the original implementations from the above models with default settings whenever available. When the code was not available, we implemented the methods based on the descriptions in the original paper using PyTorch as backend in Python. It should be noted that we used PyTorch to rewrite the source code for DeepSynergy and DTF, and employed DrugCom in MATLAB. Supplementary Tables

S1 summarizes our methods and the comparison methods. Supplementary Tables S2 shows the statistics information about hyperparameters in ConvNTC and comparison methods.

Table S1: Summarization of ConvNTC, MCTD and other comparison methods..

| Methods | Tensor factorization | Auxilary information | Linear relation | Nonlinear relation |
|---|---|---|---|---|
| **ConvNTC (ours)** | ✓ | ✓ | ✓ | ✓ |
| **MCTD (ours)** | ✓ | ✓ | ✓ | −− |
| CP | ✓ | −− | ✓ | −− |
| TFAI | ✓ | ✓ | ✓ | −− |
| DrugCom | ✓ | ✓ | ✓ | −− |
| TDRC | ✓ | ✓ | ✓ | −− |
| CTF | ✓ | ✓ | ✓ | −− |
| DeepSynergy | −− | ✓ | −− | ✓ |
| Costco | −− | −− | −− | ✓ |
| DTF | ✓ | −− | ✓ | ✓ |
| GraphTF | −− | ✓ | −− | ✓ |
| CTF-DDI | ✓ | ✓ | ✓ | ✓ |

Table S2: Hyper-parameter settings of all methods.

| Methods | $r$ | $\mu$ | $\eta$ | $\alpha$ | $\beta$ | $\lambda$ | $lr$ | $batch\_size$ | $epoch$ | $n_C$ | $\gamma$ |
|---|---|---|---|---|---|---|---|---|---|---|---|
| **ConvNTC_1** | 57 | 0.75 | 0.125 | 0.25 | 0.25 | 0.001 | 0.0001 | 256 | 500 | 114 | 0.5 |
| **ConvNTC_2** | 123 | 0.25 | 2 | 0.125 | 0.125 | 0.001 | 0.00001 | 512 | 500 | 246 | 0.8 |
| **MCTD_1** | 57 | 0.75 | 0.125 | 0.25 | 0.25 | 0.001 | −− | −− | −− | −− | −− |
| **MCTD_2** | 123 | 0.25 | 2 | 0.125 | 0.125 | 0.001 | −− | −− | −− | −− | −− |
| CP | 57/123 | −− | −− | −− | −− | −− | −− | −− | −− | −− | −− |
| TFAI | 57/123 | −− | −− | 2 | 0.125 | 0.001 | −− | −− | −− | −− | −− |
| DrugCom | 57/123 | −− | −− | 0.1 | 0.1 | −− | −− | −− | −− | −− | −− |
| TDRC | 57/123 | −− | −− | 2 | 0.125 | 0.001 | −− | −− | −− | −− | −− |
| CTF | 57/123 | 0.5 | 0.2 | 0.5 | 0.5 | 0.5 | −− | −− | −− | −− | −− |
| DeepSynergy | −− | −− | −− | −− | −− | −− | 0.00001 | 64 | 1000 | −− | −− |
| Costco | 57/123 | −− | −− | −− | −− | −− | 0.0001 /0.00001 | 256/512 | 500 | 114 | −− |
| DTF | 57/1000 | −− | −− | −− | −− | −− | 0.00001 | 128 | 1000 | −− | −− |
| GraphTF | 128/128 | −− | −− | −− | −− | −− | 0.001 | full-batch | 300 | −− | −− |
| CTF-DDI | 57/123 | 0.5 | 0.2 | 0.5 | 0.5 | 0.5 | 0.0001 | 1000 | 300 | −− | −− |

[1] ConvNTC_1 and MCTD_1 are our model applied to Dataset1, ConvNTC_2 and MCTD_2 is our model applied on Dataset2.

[2] The $r$ value in GraphTF denotes the dimension of the learned features from graph attention mechanism.

[3] The $r$ value for all baselines is $value1/value2$, where $value1$ is the rank used on Dataset1, $value2$ is the rank used on Dataset2. It should be noted that the rank for Dataset2 is 1000 in DTF (same as the setting of original paper) since Dataset2 was also used in the original paper of DTF.

### 4 Supplementary material 4: Parameters analysis

We analyze how hyperparameters influence the performance of ConvNTC. Considering that ConvNTC as a two-stage model comprising the MCTD and NTD modules, we initially determine the optimal parameter combination for the six parameters within the MCTD module to streamline the search for the overall optimal configuration. Subsequently, with these six parameters held constant, we identify the optimal combination for the remaining parameters in the NTD module.

#### 4.1 MCTD module

MCTD module has six hyperparameters:

- $r$ is the rank of the reconstructed tensor;
- $\mu, \eta$ control the contributions of miRNA and disease similarity, respectively;
- $\alpha, \beta$ are the Hessian regularization coefficients of factors $M$ and $D$, respectively;
- $\lambda$ is the $L_{2,1}$ regularization coefficient.

Compared to the other five parameters, $\lambda$ has less influence on performance. We empirically set $\lambda = 0.001$ and analyze the influence of the remaining parameters on performance of MCTD as follows:

- First, we set $r = 30$ for Dataset1, $r = 32$ for Dataset2, $\lambda = 0.001$. The ranges of $\mu, \eta, \alpha, \beta$ all in $[2^{-3}, 2^{-2}, 2^{-1}, 0.75, 2^0, 2^1]$, and we evaluated the performance of MCTD module by using grid search of these parameters on both Dataset1 and Dataset2. As a result, MCTD achieved the best performance on Dataset1 when $\mu = 0.75, \eta = 0.125, \alpha = 0.25, \beta = 0.25$, and on Dataset2 when $\mu = 0.25, \eta = 2, \alpha = 0.125, \beta = 0.125$.
- Next, we set $\mu = 0.75, \eta = 0.125, \alpha = 0.25, \beta = 0.25$ for Dataset1 and $\mu = 0.25, \eta = 2, \alpha = 0.125, \beta = 0.125$ for Dataset2, $r$ in the range of $[30, ..., 127, 128]$ on Dataset1 and $[32, ..., 127, 128]$ on Dataset2, and we evaluated $r$ using a grid search on both Dataset1 and Dataset2. As a result, MCTD achieved the best performance on Dataset1 when $r = 57$, and on Dataset2 when $r = 123$.

#### 4.2 NTD module

NTD module has five hyperparameters:

- Traditional hyperparameters of deep learning are $lr$ (denoting learning rate), $batch\_size$ and $epoch$;
- $n_C$ is the number of channels in convolutional encoder;
- $\gamma$ is the parameter to balance the weight of factor and index embeddings.

Since we designed an early stopping strategy during model training to prevent ConvNTC from overfitting, the epoch is empirically set to 500. Based on the settings of $r = 57, \mu = 0.75, \eta = 0.125, \alpha = 0.25, \beta = 0.25, \lambda = 0.001, epoch = 500$ on Dataset1 and $r = 123, \mu = 0.25, \eta = 2, \alpha = 0.125, \beta = 0.125, \lambda = 0.001, epoch = 500$ on Dataset2, we analyze the influence of the other four parameters on performance of NTD as follows:

- First, we set $\gamma = 0.5$, $lr$ in the range of $[0.001, 0.0001, 0.00001]$, $batch\_size$ in the ranges of $[256, 512, 1024]$ and $n_C$ in the range of $[\frac{1}{2}r, r, 2r]$, and we evaluated the performance of NTD module by using grid search of $lr, batch\_size, n_C$ on both Dataset1 and Dataset2. As a result, NTD achieved the best performance on Dataset1 when $lr = 0.0001$, $batch\_size = 256$, $n_C = 2r$, and Dataset2 when $lr = 0.00001$, $batch\_size = 512$, $n_C = 2r$.

- Finally, we set $n_C = 2r, lr = 0.0001, batch\_size = 256$ on Dataset1 and $n_C = 2r, lr = 0.0001, batch\_size = 256$ on Dataset2, the ranges of $\gamma$ in $[0, 0.1, 0.2, 0.3, 0.4, 0.5, 0.6, 0.7, 0.8, 0.9, 1]$, and we evaluated the performance of NTD module by using grid search of $\gamma$ on both Dataset1 and Dataset2. As a result, NTD achieved the best performance on Dataset1 when $\gamma = 0.5$, and on Dataset2 when $\gamma = 0.8$.

In conclusion, the best hyperparameters setting of ConvNTC on Dataset1 is: $r = 57, \mu = 0.75, \eta = 0.125, \alpha = 0.25, \beta = 0.25, \lambda = 0.001, n_C = 2r, lr = 0.0001, batch\_size = 256, epoch = 500, \gamma = 0.5$. The best hyperparameter setting of ConvNTC on Dataset2 is: $r = 57, \mu = 0.25, \eta = 2, \alpha = 0.125, \beta = 0.125, \lambda = 0.001, n_C = 2r, lr = 0.00001, batch\_size = 512, epoch = 500, \gamma = 0.8$. Figure S1 intuitively showd the impact of r and $\gamma$ values on the model performance across Dataset1 and Dataset2.

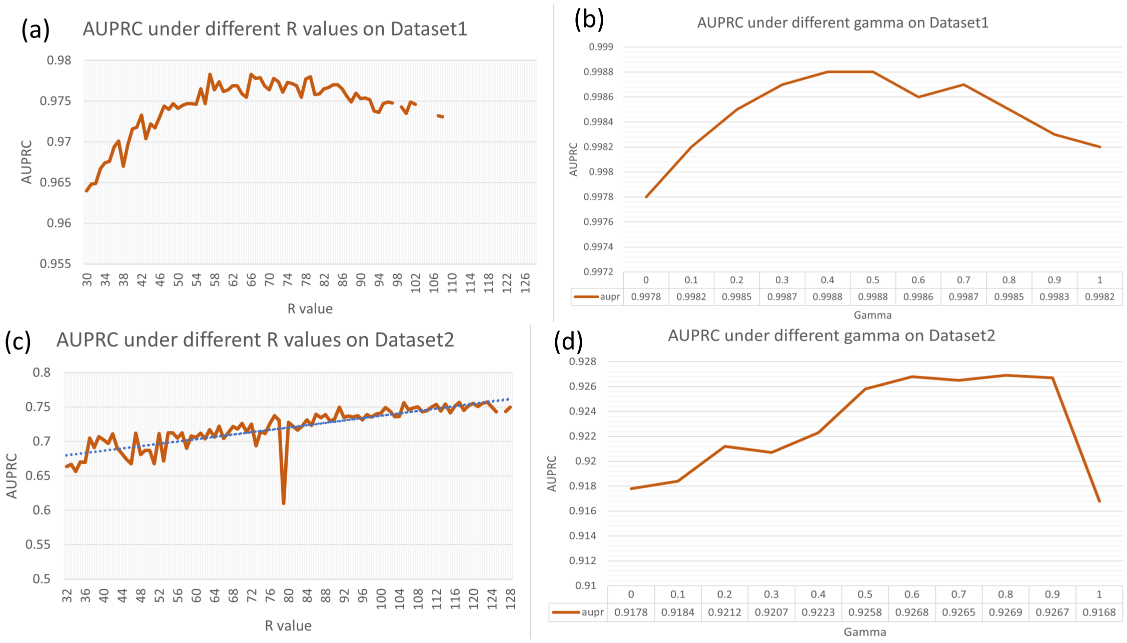

Figure S1: The impact of r and $\gamma$ values on the model performance across Dataset1 and Dataset2.

### 5 Supplementary material 5: Ablation study

ConvNTC consists of five parts, i.e., MCTD, index embedding, factor embedding, convolutional encoder and FastKAN predictor affecting on its performance. To assess the impact of these parts, we first systematically designed five variants: ConvNTC-nfact (i.e., $\gamma = 0$, no factor embedding), ConvNTC-nind (i.e., $\gamma = 1$, no index embedding), ConvNTC-nconv (i.e., no convolutional encoder), ConvNTC-mlp (i.e. FastKAN replaced with MLP), and ConvNTC-nntd (i.e., MCTD module, no NTD module). Summarization was shown in Table S3 and results are shown in Table S4.

From the results in Table S4, we found that ConvNTC achieved the best performance on both datasets using all five parts and ConvNTC-mlp underperformed across all datasets, which underscored the effectiveness of our predictor constructed using the KAN network. Specifically, on Dataset2, the performance of ConvNTC-mlp was no different from random guessing. We hypothesize that this is due to the significant class imbalance, where the positive samples number only 4,162 compared to 41,942 negative samples. This imbalance likely caused ConvNTC-mlp to overfit to the negative class, resulting in predictions that were almost entirely zeros. When calculating evaluation metrics, we employed a threshold based on the predicted scores to distinguish between positive and negative samples. Samples with scores below the threshold were classified as negative, while those equal to or above the threshold were classified as positive. Consequently, when the threshold was set to zero, all samples were classified as positive, yielding a AUROC and recall score of 0.5 and 1 respectively.

In addition, ConvNTC-nconv and ConvNTC-nntd also have a clear impression of model performance, which indicated that the entire nonlinear relationship learning module (i.e., NTD) is able to significantly enhance the performance of ConvNTC. Moreover, convolutional encoder and FastKAN predictor are indispensable parts of NTD, playing a crucial role in capturing the complex nonlinear charateristics. Together, the results clearly demonstrated the advantages of collaborative learning of multilinear and nonlinear modules of ConvNTC.

Table S3: Summarization of ConvNTC and its variants.

| Variants | MCTD | Index embedding | Factor emdedding | Convolutional encoder | Predictor |
| --- | --- | --- | --- | --- | --- |
| ConvNTC-nind | ✓ | −− | ✓ | ✓ | $FastKAN$ |
| ConvNTC-nfact | −− | ✓ | ✓ | ✓ | $FastKAN$ |
| ConvNTC-nconv | ✓ | ✓ | ✓ | −− | $FastKAN$ |
| ConvNTC-mlp | ✓ | ✓ | ✓ | ✓ | $MLP$ |
| ConvNTC-nntd | ✓ | − | −− | −− | −− |
| **ConvNTC** | ✓ | ✓ | ✓ | ✓ | $FastKAN$ |

[1] MCTD is a part of multi-linear relationship learning module; Embedding layer (including Index embedding and Factor embedding), Convolutional encoder and predictor is three parts of the nonlinear relationship learning module.

[2] ConvNTC-nind denotes we only use the predicting score derived from factor embedding to evaluate the impact of index embedding ($\gamma = 1$).

[3] ConvNTC-nfact denotes we only use the predicting score derived from index embedding to evaluate the impact of factor embedding ($\gamma = 0$).

[4] ConvNTC-nconv denotes the convolutional encoder is removed to assess the importance of it.

[5] ConvNTC-mlp denotes the FastKAN is replaced by MLP to evaluate the effectiveness of FastKAN.

[6] ConvNTC-nntd (i.e. MCTD module) denotes only the MCTD module is retained to obtain the predicting score to evaluate the impact of the whole nonlinear relationship learning moduled.

Table S4: Comparison results of ablation study on Dataset1 and Dataset2. The bolded items indicate the optimal performance.

| Methods | AUPRC | AUROC | F1 | Accuracy | Recall | Specificity | Precision | Paired t-test |
|---|---|---|---|---|---|---|---|---|
| | | | | Dataset1 | | | | |
| ConvNTC-mlp | 0.9546 | 0.9151 | 0.9297 | 0.9030 | 0.9836 | 0.8223 | 0.9024 | 0.00539 |
| ConvNTC-nconv | 0.9731 | 0.9735 | 0.9290 | 0.9284 | 0.9365 | 0.9204 | 0.9218 | 0.00035 |
| ConvNTC-nntd | 0.9780 | 0.9706 | 0.9234 | 0.9263 | 0.8905 | 0.9621 | 0.9594 | 0.00523 |
| ConvNTC-nind | 0.9978 | 0.9980 | 0.9840 | 0.9839 | 0.9851 | 0.9828 | 0.9829 | 0.00379 |
| ConvNTC-nfact | 0.9982 | 0.9986 | 0.9889 | 0.9888 | 0.9925 | 0.9851 | 0.9852 | 0.00756 |
| **ConvNTC** | **0.9988** | **0.9991** | **0.9926** | **0.9926** | **0.9946** | **0.9905** | **0.9906** | |
| | | | | Dataset2 | | | | |
| ConvNTC-nconv | 0.4249 | 0.8525 | 0.4367 | 0.8844 | 0.4962 | 0.9229 | 0.3957 | 0.00670 |
| ConvNTC-mlp | 0.5451 | 0.5000 | 0.1656 | 0.0903 | 1.0000 | 0.0000 | 0.0903 | 0.00660 |
| ConvNTC-nntd | 0.7563 | 0.9574 | 0.7155 | 0.9476 | 0.7298 | 0.9692 | 0.7033 | 0.01070 |
| ConvNTC-nind | 0.9168 | 0.9827 | 0.8790 | 0.9786 | 0.8607 | 0.9903 | 0.8987 | 0.00980 |
| ConvNTC-nfact | 0.9178 | 0.9835 | 0.8853 | 0.9798 | 0.8618 | 0.9915 | 0.9105 | 0.02790 |
| **ConvNTC** | **0.9269** | **0.9853** | **0.8901** | **0.9806** | **0.8691** | **0.9917** | **0.9126** | |

### 6 Supplementary material 6: Comparative experiments

#### 6.1 Robustness analysis

Figure S2 depicted the AUPRC values of all methods in different negative sample sizes.

Figure S3 depicted the AUROC values of all methods in different training ratios.

#### 6.2 Generalizability analysis

To assess the generalizability of ConvNTC, we designed a new experiment to investigate whether ConvNTC can predict synergy status for new unseen data. Specifically, we conduct five-fold cross-validation in five times on entities (i.e., diseases or cell lines) rather than triple associations on two datasets, and the averaged results are reported. Detailed values of all metrics are shown in Table S5. In addition to the observations mentioned in the main paper, it was worth mentioning that on Dataset1, the AUROC, recall, and specificity

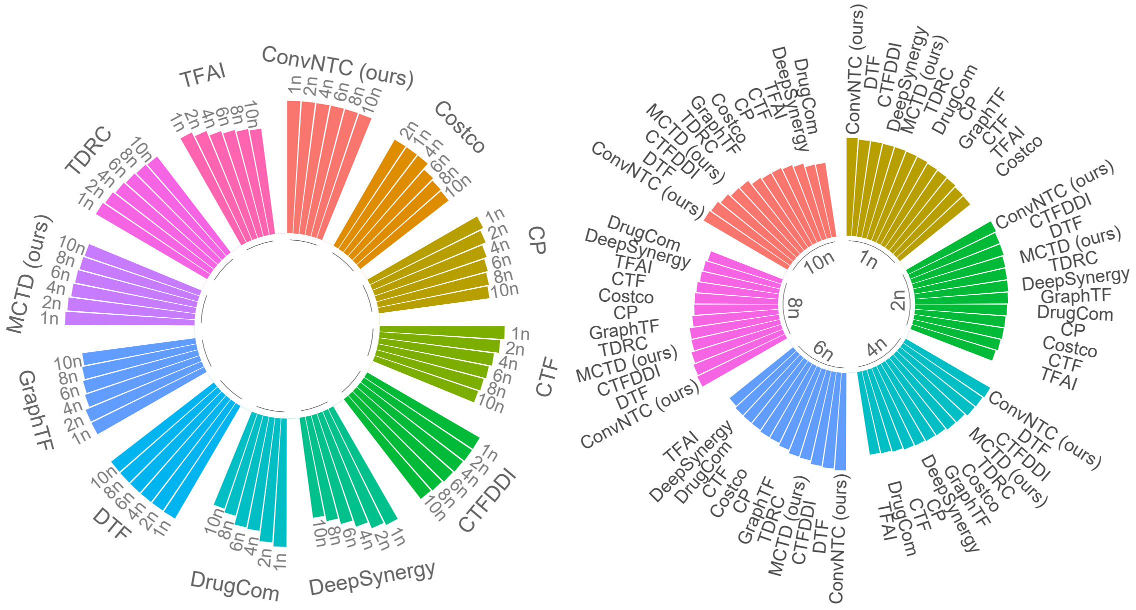

Figure S2: The AUPRC values of all methods in different negative sample sizes. The right panel displayed results grouped by different methods, while the left panel presented results grouped by varying negative sample sizes.

of the four linear methods—CP, TDRC, TFAI, and CTF—were 0.5, 1, and 0, respectively. This indicated that their performance in predicting miRNAs associated with novel diseases is equivalent to random guessing. We speculated that the reason for this result is the same as that on Dataset2 with ConvNTC-mlp in the ablation experiment. The class imbalance (with too few positive samples) caused the model to fit the negative samples, leading to prediction outputs that were almost entirely zeros. This also indicated that the generalization capability of these models is relatively poor.

### 7 Supplementary material 7: Case Studies

#### 7.1 Predicting the novel miRNA-miRNA-disease triplets

##### 7.1.1 TAM 2.0

TAM 2.0 is a web tool for miRNA set enrichment analysis, which groups miRNAs into six categories of miRNA sets: miRNA-family sets, miRNA cluster sets, miRNA-disease, miRNA-function sets, miRNA-TF sets and tissue specificity sets. Given a miRNA set, TAM 2.0 will enrich all miRNAs in this set from the above six groups. First, the results of enrichment analysis for each miRNA pair (containing two miRNAs) in six categories was obtained by using TAM. Then, based on the condition that each term contains at least

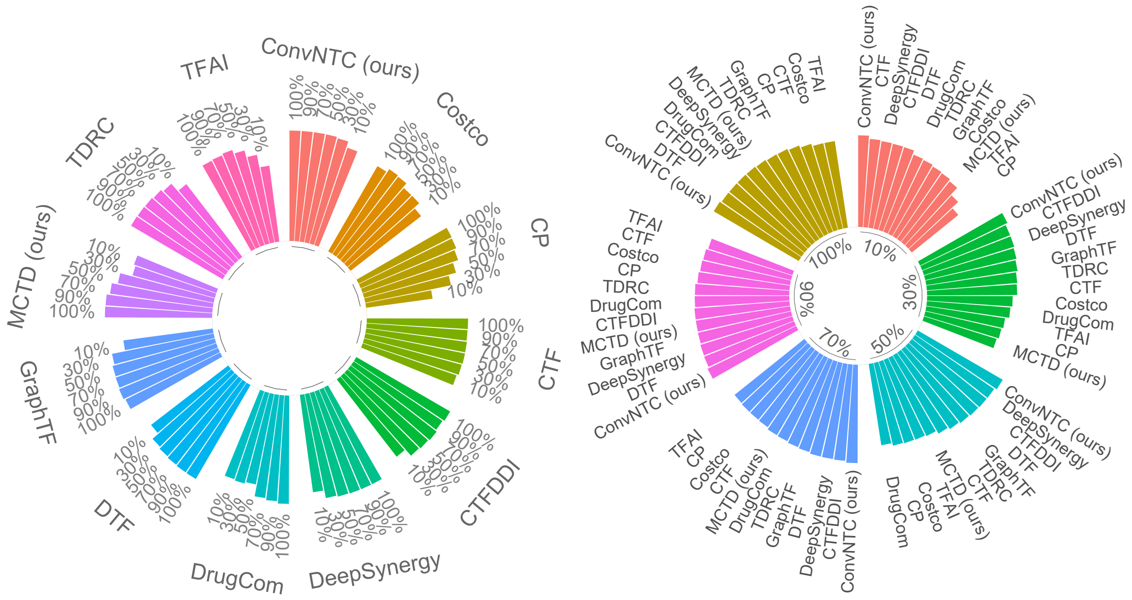

Figure S3: The AUROC values of all methods in different training ratios. The right panel displayed results grouped by different methods, while the left panel presented results grouped by varying training ratios.

two miRNAs and $p\_value < 0.05$, we believed that this term are co-enriched by two miRNAs in each miRNA pair. Finally, the average $-log10(p\_value)$ of all enriched terms by category was taken as the enrichment score of each miRNA pair in each category.

###### 7.1.2 Association network of the top-10 triple predictions

Figure S5 displays the association network of the top-10 triple predictions, where we observe that most miRNAs act on diseases in the form of pairs (or even modules).

###### 7.1.3 Survival analysis

To further verify the biological significance of predictions, we performed survival analysis on top-10 predicted miRNA pairs using the preprocessed miRNA expression data and clinical data for breast cancer from TCGA, which were obtained through the R package UCSCXenaTools (Goldman et al., 2020). The primary metric used in our analysis was the risk score, constructed via multivariable Cox proportional hazards regression, based on both miRNA expression and clinical data. Kaplan-Meier survival analysis was then employed to assess the prognostic significance of these miRNA pairs.

Figure S6 depicted the survival curves and pearson correlation of miRNA expression for three novel miRNA pairs, i.e., (hsa-mir-125a, hsa-mir-99b), (hsa-mir-99a, hsa-mir-99b) and (hsa-mir-181a-2, hsa-mir-181d), that significantly explains the survival rate of breast cancer patients.

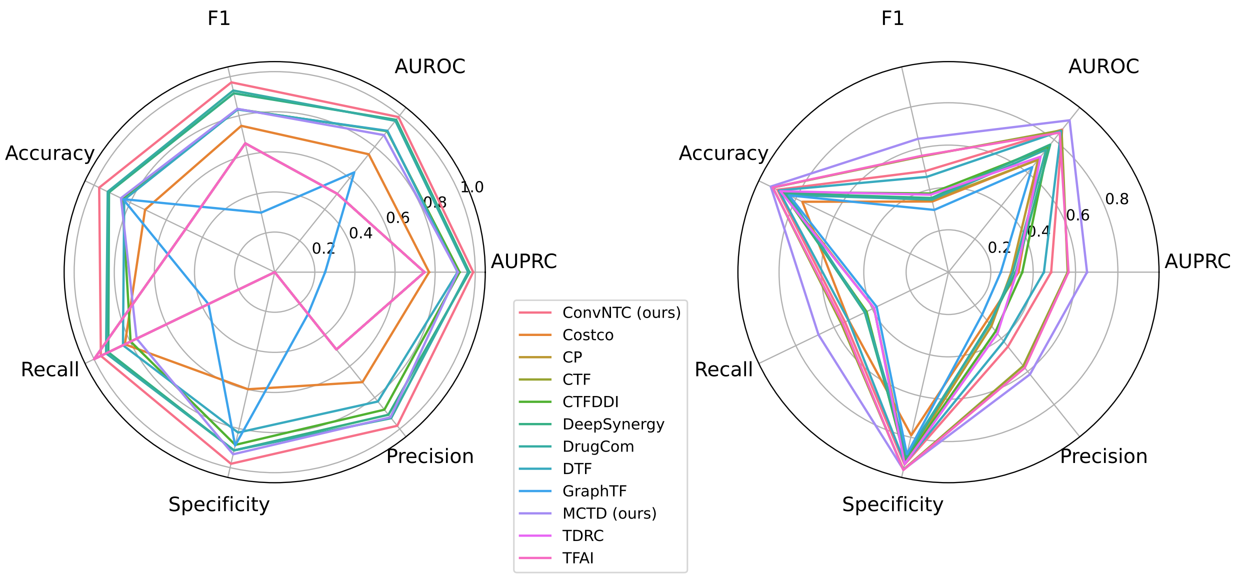

Figure S4: Comparison results of all methods in generalizability analysis experiments on two datasets (Dataset1 on right panel and Dataset2 on left panel).

###### 7.1.4 Enrichment analysis

The enrichment analysis was performed on these three pairs. For each pair, its gene set is composed of common target genes of each miRNA. The target genes of each pre-miRNA is defined as the union of target genes between its mature miRNAs from miRTarbase v9.0 (Huang et al., 2022). We first used the R package ‘clusterProfiler’ (Yu et al., 2012) and ’ReactomePA’ (Yu and He, 2016) to explore whether gene sets are involved in significant GO-BP terms, Wikipathways and Reactome pathways ($p\_value < 0.05$). Then, we combined top-5 enriched terms and pathways in GO-BP, Wikipathways and Reactome respectively, and show them in Figure 2 of main paper.

Figure 2 depicted the enriched terms and pathways that significantly impact breast cancer development and progression. Key processes like ’lysosome organization‘ and ’vacuole organization‘ help cancer cells survive under stress by regulating autophagy and cellular homeostasis. ’Chromosome segregation‘ and ’mitotic phase regulation‘ are critical for maintaining genomic stability, with errors leading to increased cancer cell proliferation. The ’Breast cancer pathway‘ is central, directly involving hormonal signaling, cell growth, and survival mechanisms specific to breast cancer. Other pathways like ’PI3K/Akt/mTOR signaling‘ drive tumor growth by enhancing cell survival and metabolism, while ’ATM signaling‘ is crucial for DNA damage repair, with mutations increasing the risk of breast cancer. Collectively, these enriched terms and pathways reveal how metabolic, genetic, and signaling dysregulation fuel breast cancer progression.

Table S5: Comparison results of all methods based across both two datasets at levels of new cell lines and new diseases, respectively.

| | Methods | AUPRC | AUROC | F1 | Accuracy | Recall | Specificity | Precision | Paired t-test |
|---|---|---|---|---|---|---|---|---|---|
| | | | | | Dataset1 | | | | |
| Linear | CTF | 0.7468 | 0.5000 | 0.6590 | 0.4936 | **1.0000** | **0.0000** | 0.4936 | 0.01149 |
| | TDRC | 0.7468 | 0.5000 | 0.6590 | 0.4936 | **1.0000** | **0.0000** | 0.4936 | 0.01149 |
| | TFAI | 0.7468 | 0.5000 | 0.6590 | 0.4936 | **1.0000** | **0.0000** | 0.4936 | 0.01149 |
| | CP | 0.7468 | 0.5000 | 0.6590 | 0.4936 | **1.0000** | **0.0000** | 0.4936 | 0.01149 |
| | MCTD (ours) | 0.9136 | 0.8742 | 0.8356 | 0.8509 | 0.7642 | 0.9316 | 0.9256 | 0.00167 |
| | DrugCom | 0.9645 | 0.9651 | 0.9291 | 0.9243 | 0.9334 | 0.9114 | 0.9315 | 0.00053 |
| Nonlinear | GraphTF | 0.2516 | 0.6366 | 0.3045 | 0.8393 | 0.3668 | 0.8869 | 0.2670 | 0.00403 |
| | Costco | 0.7699 | 0.7521 | 0.7485 | 0.7175 | 0.8294 | 0.5989 | 0.7017 | 0.00012 |
| | DTF | 0.9119 | 0.9017 | 0.8319 | 0.8330 | 0.8399 | 0.8207 | 0.8251 | 0.00005 |
| | CTFDDI | 0.9220 | 0.8992 | 0.8337 | 0.8443 | 0.7975 | 0.8835 | 0.8772 | 0.00010 |
| | DeepSynergy | 0.9712 | 0.9716 | 0.9150 | 0.9171 | 0.9207 | 0.9120 | 0.9095 | 0.00125 |
| | **ConvNTC (ours)** | **0.9898** | **0.9895** | **0.9709** | **0.9715** | **0.9622** | **0.9801** | **0.9799** | |
| | | | | | Dataset2 | | | | |
| Linear | CP | 0.2967 | 0.6795 | 0.3487 | 0.8655 | 0.3905 | 0.9120 | 0.3255 | 0.00346 |
| | DrugCom | 0.3042 | 0.7408 | 0.3517 | 0.8405 | 0.4359 | 0.8802 | 0.3134 | 0.00065 |
| | TDRC | 0.3225 | 0.6976 | 0.3752 | 0.8813 | 0.3896 | 0.9296 | 0.3708 | 0.00875 |
| | CTF | 0.5628 | 0.8567 | 0.5608 | 0.9207 | 0.5611 | 0.9561 | 0.5665 | 0.02724 |
| | TFAI | 0.5670 | 0.8428 | 0.5639 | 0.9218 | 0.5541 | 0.9588 | 0.5779 | 0.04732 |
| | **MCTD (ours)** | **0.6544** | **0.9173** | **0.6459** | **0.9315** | **0.6813** | **0.9563** | **0.6173** | 0.00502 |
| Nonlinear | GraphTF | 0.2472 | 0.6344 | 0.3015 | 0.8354 | 0.3766 | 0.8822 | 0.2555 | 0.00136 |
| | DeepSynergy | 0.3177 | 0.7705 | 0.3592 | 0.8573 | 0.4386 | 0.8981 | 0.3098 | 0.00181 |
| | Costco | 0.3308 | 0.6808 | 0.3421 | 0.7662 | 0.5234 | 0.7905 | 0.2924 | 0.00054 |
| | CTFDDI | 0.3485 | 0.7538 | 0.3812 | 0.8711 | 0.4294 | 0.9140 | 0.3573 | 0.00249 |
| | DTF | 0.4507 | 0.8527 | 0.4614 | 0.8892 | 0.5244 | 0.9249 | 0.4184 | 0.00498 |
| | ConvNTC (ours) | 0.4849 | 0.8615 | 0.4896 | 0.8972 | 0.5423 | 0.9319 | 0.4501 | |

#### 7.2 Predicting novel synergistic drug pairs

Table 3 in main paper outlines the top-10 predicted drug combinations for the A375 melanoma cancer cell line, 9 out of 10 supported by scientific evidence. The combinations focus on targeting various cellular processes, aiming to enhance the efficacy of cancer treatments by leveraging the complementary mechanisms of different drugs:

1. Erlotinib (EGFR inhibitor) is paired with several drugs that inhibit different pathways:
   - Talazoparib (PARP inhibitor) is known to induce DNA damage in cancer cells, and when combined with Erlotinib, it can exploit the weakened DNA repair mechanism in cancer cells while also blocking EGFR-driven growth, potentially leading to enhanced cell death.
   - Lenvatinib targets multiple tyrosine kinases involved in angiogenesis and cell proliferation. Its combination with Erlotinib could synergistically inhibit both cancer cell signaling and blood vessel formation, which is critical for tumor growth.
   - Everolimus (mTOR inhibitor) affects the mTOR pathway, which is central to cancer cell metabolism

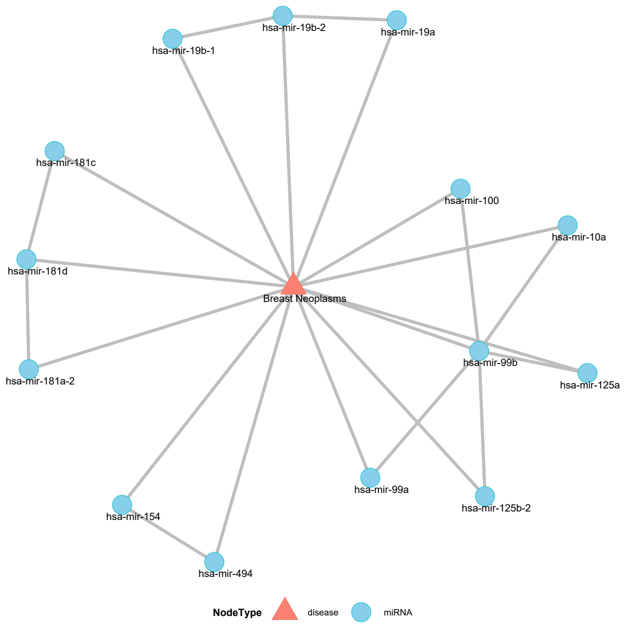

Figure S5: The association network of the top-10 triple predictions for Breast Neoplasms. Noting that 'known' means the projected pairwise associations observed in the original data, and 'confirmed' means the unknown associations confirmed by existing databases and TAM 2.0.

and survival. This combination may prevent both growth signaling and metabolic support for the cancer cells.

- Copanlisib is a PI3K inhibitor, targeting another critical survival pathway in cancer cells. Combined with Erlotinib, this could lead to the disruption of both EGFR and PI3K pathways, limiting tumor cell proliferation and survival.

2. Abemaciclib + Erlotinib:

    - Abemaciclib is a CDK4/6 inhibitor that regulates the cell cycle, while Erlotinib targets growth signaling through EGFR inhibition. The combination may be effective in halting cell division and simultaneously inhibiting growth signals, making it more difficult for the cancer cells to continue proliferating.

3. Erlotinib + Crizotinib or Pazopanib:

    - Crizotinib and Pazopanib both target different tyrosine kinases involved in tumor growth and metastasis. Combined with Erlotinib, these drugs offer a multi-faceted approach to shutting down critical pathways that cancer cells use for survival and spread.

Overall, these combinations emphasize a multi-targeted approach by inhibiting growth signals (EGFR,

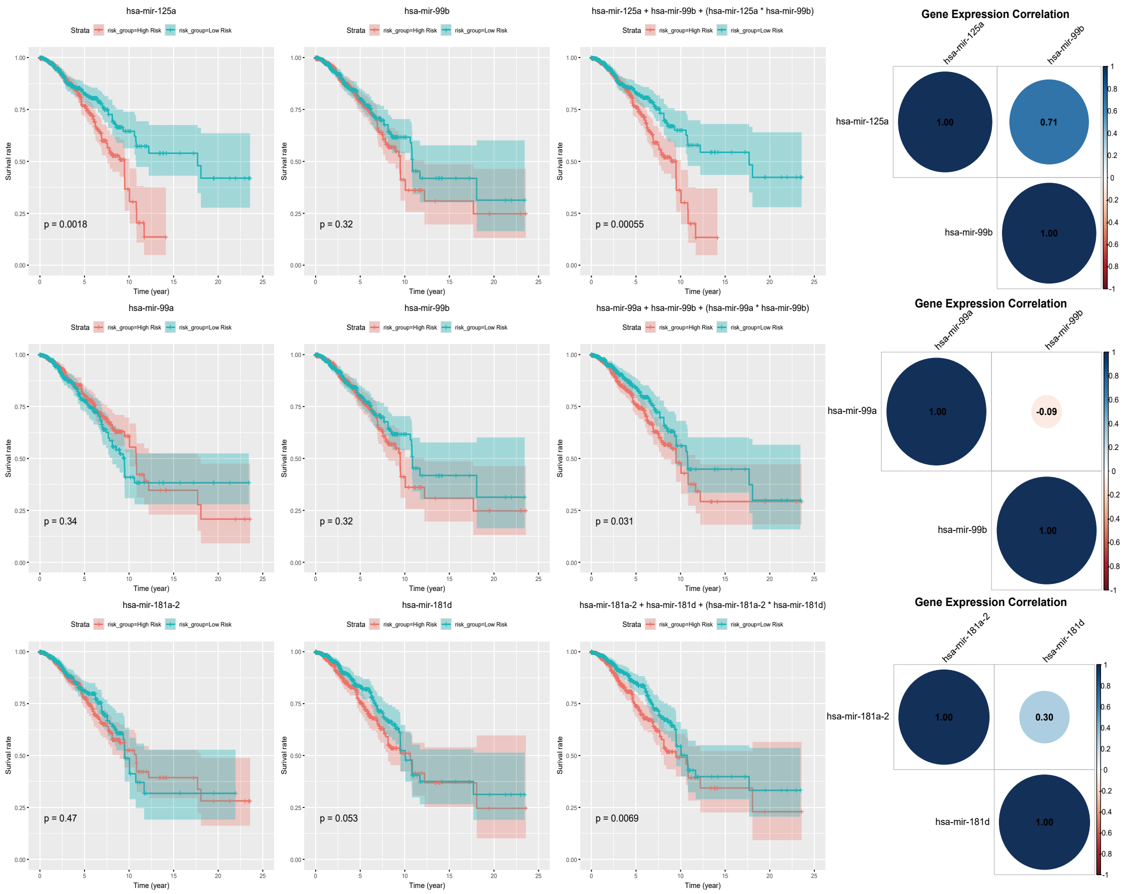

Figure S6: The survival curves and expression correlation of miRNA pairs (hsa-mir-125a, hsa-mir-99b), (hsa-mir-99a, hsa-mir-99b) and (hsa-mir-181a-2, hsa-mir-181d) for Breast Neoplasms. Specifically, for the pair (hsa-mir-125a, hsa-mir-99b), the combined analysis revealed a higher pronounced statistical distinction ($p\_value < 0.5$) between high-risk and low-risk groups compared to the individual analyses of hsa-mir-125a and hsa-mir-99b. Similar phenomena occurred in the other two pairs. Moreover, the expressions of pairs (hsa-mir-125a, hsa-mir-99b) and (hsa-mir-181a-2, hsa-mir-181d) were positively correlated, whereas the pair (hsa-mir-99a, hsa-mir-99b) was negatively correlated.

PI3K, mTOR), promoting DNA damage (PARP inhibition), and disrupting the cell cycle (CDK4/6 inhibition). These combinations aim to block multiple cancer survival mechanisms simultaneously, offering a potentially more effective therapeutic strategy for treating A375 melanoma cells.

### References

H. Chen and J. Li. Drugcom: Synergistic discovery of drug combinations using tensor decomposition. In *2018 IEEE International Conference on Data Mining (ICDM)*, pages 899–904. IEEE, 2018.

M. J. Goldman, B. Craft, M. Hastie, K. Repečka, F. McDade, A. Kamath, A. Banerjee, Y. Luo, D. Rogers, A. N. Brooks, et al. Visualizing and interpreting cancer genomics data via the xena platform. *Nature biotechnology*, 38(6):675–678, 2020.

G. Han, L. Peng, A. Ding, Y. Zhang, and X. Lin. Ctf-ddi: Constrained tensor factorization for drug–drug interactions prediction. *Future Generation Computer Systems*, 161:26–34, 2024.

F. Huang, X. Yue, Z. Xiong, Z. Yu, S. Liu, and W. Zhang. Tensor decomposition with relational constraints for predicting multiple types of microrna-disease associations. *Briefings in bioinformatics*, 22(3):bbaa140, 2021.

H.-Y. Huang, Y.-C.-D. Lin, S. Cui, Y. Huang, Y. Tang, J. Xu, J. Bao, Y. Li, J. Wen, H. Zuo, et al. mirtarbase update 2022: an informative resource for experimentally validated mirna–target interactions. *Nucleic acids research*, 50(D1):D222–D230, 2022.

Z. Huang, J. Shi, Y. Gao, C. Cui, S. Zhang, J. Li, Y. Zhou, and Q. Cui. Hmdd v3. 0: a database for experimentally supported human microrna–disease associations. *Nucleic acids research*, 47(D1):D1013–D1017, 2019.

T. G. Kolda and B. W. Bader. Tensor decompositions and applications. *SIAM review*, 51(3):455–500, 2009.

A. Kozomara, M. Birgaoanu, and S. Griffiths-Jones. mirbase: from microrna sequences to function. *Nucleic acids research*, 47(D1):D155–D162, 2019.

Z. Li. Kolmogorov-arnold networks are radial basis function networks. *arXiv preprint arXiv:2405.06721*, 2024.

H. Liu, Y. Li, M. Tsang, and Y. Liu. Costco: A neural tensor completion model for sparse tensors. In *Proceedings of the 25th ACM SIGKDD International Conference on Knowledge Discovery & Data Mining*, pages 324–334, 2019.

P. Liu, J. Luo, and X. Chen. mircom: tensor completion integrating multi-view information to deduce the potential disease-related mirna-mirna pairs. *IEEE/ACM Transactions on Computational Biology and Bioinformatics*, 19(3):1747–1759, 2020.

Z. Liu, Y. Wang, S. Vaidya, F. Ruehle, J. Halverson, M. Soljačić, T. Y. Hou, and M. Tegmark. Kan: Kolmogorov-arnold networks. *arXiv preprint arXiv:2404.19756*, 2024.

J. Luo, Z. Lai, C. Shen, P. Liu, and H. Shi. Graph attention mechanism-based deep tensor factorization for predicting disease-associated mirna-mirna pairs. In *2021 IEEE International Conference on Bioinformatics and Biomedicine (BIBM)*, pages 189–196. IEEE, 2021.

A. Narita, K. Hayashi, R. Tomioka, and H. Kashima. Tensor factorization using auxiliary information. *Data Mining and Knowledge Discovery*, 25:298–324, 2012.

K. Preuer, R. P. Lewis, S. Hochreiter, A. Bender, K. C. Bulusu, and G. Klambauer. Deepsynergy: predicting anti-cancer drug synergy with deep learning. *Bioinformatics*, 34(9):1538–1546, 2018.

Z. Sun, S. Huang, P. Jiang, and P. Hu. Dtf: deep tensor factorization for predicting anticancer drug synergy. *Bioinformatics*, 36(16):4483–4489, 2020.

G. Yu and Q.-Y. He. Reactomepa: an r/bioconductor package for reactome pathway analysis and visualization. *Molecular BioSystems*, 12(2):477–479, 2016.

G. Yu, L.-G. Wang, Y. Han, and Q.-Y. He. clusterprofiler: an r package for comparing biological themes among gene clusters. *Omics: a journal of integrative biology*, 16(5):284–287, 2012.
